# Hybrid hydrogel-extracellular matrix scaffolds identify distinct ligand and mechanical signatures in cardiac aging

**DOI:** 10.1101/2023.10.06.561048

**Authors:** Avery Rui Sun, Md. Faris H. Ramli, Xingyu Shen, Dixiao Chen, Roger S. Foo, Jin Zhu, Matthew Ackers-Johnson, Jennifer L. Young

**Affiliations:** Mechanobiology Institute (MBI), National University of Singapore, 5A Engineering Drive 1, 117411, Singapore; Department of Biomedical Engineering, College of Design and Engineering, National University of Singapore, 4 Engineering Drive 3, 117583, Singapore; Cardiovascular-Metabolic Diseases Translational Research Programme, Yong Loo Lin School of Medicine, National University of Singapore, Singapore; Cardiovascular Research Institute, National University Healthcare Systems, Singapore

**Keywords:** Cardiac fibroblast, myofibroblast, stiffness, tunable, ligand presentation, mechanotransduction

## Abstract

Extracellular matrix (ECM) remodeling of cardiac tissue is a key contributor to age-related cardiovascular disease and dysfunction. Aberrant secretion, structural perturbations, and degradation of specific ECM components lead to significant alterations in ECM properties that disrupt healthy cell and tissue homeostasis. These changes in ECM are multifaceted, as alterations in ligand presentation, including both biochemical and architectural aspects, are often accompanied by stiffness changes, clouding our understanding of how and which ECM properties contribute to a dysfunctional state. To identify the specific roles of these interconnected ECM cues and elucidate their mechanistic regulation in cellular function, we developed a material system that can independently present these two distinct matrix properties, i.e., ligand presentation and stiffness, to cultured cells *in vitro*. We describe a decellularized ECM-synthetic hydrogel hybrid scaffold that maintains native matrix composition and organization of young or aged murine cardiac tissue with independently tunable scaffold mechanics that mimic young or aged tissue stiffness. Seeding these scaffolds with primary cardiac fibroblasts (CFs) from young or aged mice, we identify distinct age- and ECM-dependent mechanisms of CF activation. Importantly, we show that ligand presentation of young ECM can outweigh profibrotic stiffness cues typically present in aged ECM in maintaining or driving CF quiescence, thereby highlighting the unique roles of ECM in aging. Ultimately, these tunable scaffolds can enable the discovery of specific ECM targets to prevent aging dysfunction and promote rejuvenation.

**DECIPHER**: **DEC**ellularized ***I****n Situ* **P**olyacrylamide **H**ydrogel-**E**CM hyb**R**id

## Introduction

Alterations in tissue properties during aging have been recognized as key drivers of organ dysfunction and disease. While much remains unknown about what happens to the extracellular matrix (ECM) at the micro- and nano-length scales during these large-scale remodeling events, it is widely acknowledged that mechanics, organization, and composition of the ECM vary with age. For instance, it has been shown that the cardiac muscle stiffens^1^, while ECM composition and organization undergoes age-dependent modifications^2–4^. Cardiac fibroblasts (CFs) are the resident cells largely responsible for remodeling of the heart tissue and are known to be mechanosensitive from both *in vitro* and *in vivo* studies^5,6^.

In healthy tissue, CFs largely remain in a quiescent state, but external stimuli, including biochemical, structural, and mechanical cues^5–7^, are able to activate quiescent CFs, leading to their differentiation into a proto-myofibroblast phenotype and subsequently into a mature myofibroblast phenotype when these stimuli are impactful and persistent^6,8^. The process of CF activation and proper myofibroblast maturation are essential for ECM deposition and the maintenance of matrix homeostasis but can also lead to fibrosis and result in functional consequences^9^. This is important in aging tissues, as alterations in the ECM can be vast and multifaceted, thereby leading to the activation of CFs and subsequent aberrant tissue remodeling^6,10^.

Indeed, it has been shown that myofibroblasts are more abundant in aged vs. young hearts and directly induce changes to tissue geometry^10–12^. While *in vitro* material systems have identified individual properties of the ECM that play distinct roles in CF function, it remains a challenge to vary these properties independently. In most scaffold platforms, tuning mechanical properties will alter the ligands and/or architecture. A handful of novel material systems have been described that are capable of independent tunability^13–15^, yet the incorporation of native ECM properties is still lacking. Thus, our mechanistic understanding of the specific contributions stemming from ECM cues is currently limited. We therefore sought to develop a native ECM-based scaffold in which we could individually tune mechanics while faithfully mimicking the *in vivo* cardiac environment, both composition and architecture, allowing for the identification of ECM-specific roles in age-related CF activation.

To do this, we developed a novel approach called DECIPHER (**DEC**ellularized ***I****n Situ* **P**olyacrylamide **H**ydrogel-**E**CM hyb**R**id) that integrates polyacrylamide (PA) hydrogel-linked cardiac tissue slices and *in situ* decellularization of ECM. We demonstrate that our decellularization technique is not only able to maintain ECM composition and fiber architecture, but that samples can be tuned to exhibit young or aged stiffness, ~10 or 40 kPa, respectively, regardless of the native stiffness of the tissue pre-decellularization. Seeding young and aged CFs onto combinations of young or aged matrix with young or aged stiffness, we demonstrate distinct age-dependent mechanisms of CF activation. While we observe that the myofibroblast transition can be driven by multiple matrix cues, ECM ligand presentation can outweigh ECM mechanics, with young ECM promoting CF quiescence despite the profibrotic stiffness cues of aged ECM stiffness. These findings provide unique insight into the distinct roles of ECM in regulating age-related cardiac dysfunction, which have important implications in matrix-based treatment strategies.

## Materials and methods

### Tissue handling and preparation

Young (1-2 months) and aged (18-24 months) hearts were extracted from C57BL/6J strain mice (obtained via NUS Animal Tissue Sharing Programme under IACUC, NUS) after euthanasia and quickly transferred to sterile 1X PBS to minimize blood clot formation. The major vascular tissues superiorly attached to hearts were then carefully removed with sterile surgical instruments. Prior to slicing the heart using a vibratome (Leica VT 1200S), a 4% low melting point agarose (Invitrogen) solution was prepared in sterile 1X PBS and kept in a 37°C water bath. The heart was then carefully immersed within the agarose embedding solution and spatially adjusted such that coronal slices are obtained during slicing. The mold was put on ice to solidify the agarose, after which the agarose-embedded heart was attached to the slicing stage with super glue and loaded onto the vibratome. Following the initial trimmings of the agarose cube, the slicing speed was set to 0.18 mm/s and 150 μm slice thickness.

### Preparation of hydrogel solutions

Coverslips (ϕ=15mm) were cleaned via sonication during progressive washes with acetone, ethanol, and MilliQ water for 10 min each and then air dried. For hydrogel attachment, coverslips were treated with a UV/ozone cleaner (Bioforce ProCleaner^TM^ Plus) for 5 min followed by immersion in a solution containing 20 mL ethanol, 600 μL 10% acetic acid, and 200 μL 3-trimethoxysilylpropylmethacrylate for 5 min. Coverslips were then washed twice in ethanol and air-dried prior to placing the tissue slice and hydrogel solution onto it. The polyacrylamide (PA)-based hydrogels used in this study mimicking the young (~10 kPa) and aged (~40 kPa) cardiac tissue stiffness were labeled as “soft” and “stiff” for easy reference, with their composition detailed in Table 1. Prepared solutions of acrylamide and bis-acrylamide were pre-reacted with formaldehyde at 4 °C for 3 hours at a stoichiometric ratio of >10:1 to ensure no excess formaldehyde in the solution as previously described^16^. Prior to hydrogel formation, Irgacure 2959 (2-Hydroxy-4′-(2-hydroxyethoxy)-2-methylpropiophenone, Sigma-Aldrich) was added for UV photoinitiation. For nanoindentaion experiments, plain PA hydrogel samples were fabricated without formaldehyde. For hydrogel visualization experiments, separate samples of 10 kPa solution were made substituted with Nile blue-tagged acrylamide (Polysciences Inc. #25395-100) at 1:1000.

### Tissue stabilization

After tissue sectioning, vibratome slices were carefully transferred to a hydrophobic glass slide (DCDMS-treated) sitting in a humid condensation chamber. The agarose surrounding the cardiac tissue was gently removed, as well as any excess liquid on the cardiac slices by wicking away at the edges with a kimwipe. 21 ± 2 μL of the hydrogel solutions were then added to each 150 μm thick cardiac slice (solution volume was adjusted based on the volume of the tissue in order to fill the coverslip area) and incubated at 4°C for 60 mins in the dark. Crosslinking was then initiated using the same parameter as plain hydrogel samples (35 mW/cm^2^ for 3 min). The samples were then transferred to a 12-well plate and kept in sterile MilliQ water at 4°C prior to decellularization.

### Decellularization of PA-stabilized cardiac tissue sections

All solutions were aseptically prepared, and all experiment areas were sterilized frequently with 70% ethanol to prevent microbial contamination. Sodium deoxycholate (SDC, Sigma-Aldrich) was dissolved in MilliQ water at 5% (w/v). The PA-stabilized tissue samples were exchanged with 2 mL/well SDC solution and kept on a gentle shaker at room temperature for 2.5 days with one exchange at 24 hrs. The samples were then washed with MilliQ water and 10,000 U penicillin/streptomycin (pen/strep, Gibco) for 2 hrs. Subsequently, 2 mL of 300 kU/mL Deoxyribonuclease-I solution (DNase-I, Sigma-Aldrich, dissolved in 0.15 M NaCl supplemented with 5 mM CaCl_2_ and 1% pen/strep) was added to each sample after discarding the pen/strep and gently shaken for 3.5 days followed by a final MilliQ water wash for 2 hours. In rare cases where sample edges detached from coverslips or there was suspected microbial contamination, samples were discarded and excluded from following experiments.

### ECM compositional assays

Quantitative compositional analyses of samples were performed for DNA, sulfated glycosoaminoglycan (sGAG), and collagen content. The sample’s DNA was extracted using phenol-chloroform and kept at −20°C until applying the PicoGreen dsDNA assay kit (Invitrogen). Total sGAGs of the samples were obtained via papain (Sigma-Aldrich) extraction followed by Blyscan^TM^ sGAG assay (Biocolor). For collagen quantification, the samples were first homogenized and hydrolyzed in concentrated hydrochloric acid (HCl) at 120°C for 3 hours before being quantified using hydroxyproline assay kit (Sigma-Aldrich). A microplate reader (Promega GloMax) was used for all assays as follows: 1) dsDNA assay: fluorescence at Ex480 nm/Em520 nm; 2) sGAG assay: absorbance at 656 nm; 3) Hydroxyproline assay: absorbance at 560 nm. Duplicate technical replicates with n=3 biological replicates were measured along with standard curves for quantification, according to suppliers’ instructions.

### Immunohistochemistry (IHC)

Immunohistochemistry (IHC) of native tissue sections and DECIPHER samples was carried out for ECM components and cellular structures (antibodies and dilutions detailed in Table 2). Briefly, samples were fixed with 4% formaldehyde (Sigma-Aldrich) and rinsed with 1×PBS before blocking and permeabilizing with 2% bovine serum albumin (BSA, Sigma-Aldrich) in 0.2% Triton X-100 (Sigma-Aldrich) for 30 minutes. Samples were then incubated with primary antibodies in 2% BSA on a gentle shaker at room temperature for 3 days followed by three times of 1X PBS washing and incubation of secondary antibodies and CF® Phalloidin (Biotium) staining for 1 hour at room temperature in the dark. To label any denatured collagen, Cy3-conjugated collagen hybridizing peptide (R-CHP, 3Helix) was diluted to 5 μM and used to stain the samples after CNA35 labeling according to product instruction^17,18^. The samples were further stained with Hoechst 33342 (Invitrogen) and mounted using Fluoromount-G^TM^ (Invitrogen). For hydrogel visualization experiments using Nile blue-tagged acrylamide, samples were co-stained with CNA35.

### Scanning electron microscopy (SEM)

Scanning electron microscopy (SEM) was used to visualize the nano-architecture of PA hydrogel-stabilized cardiac tissue after decellularization (DECIPHER samples) and native cardiac tissue cryosections cut at 10 µm (Leica CM1950) from flash-frozen fresh young hearts embedded in OCT (Tissue-Tek, Sakura Finetek USA). Briefly, prior to SEM, samples were fixed with glutaraldehyde (Sigma-Aldrich) followed by dehydration using a series of ethanol exchanges of increasing concentration and dried using a Critical Point Dryer (Tousismis Autosamdri-815). The samples were coated with 8 nm platinum before being imaged with a Hitachi Regulus 8230 FE-SEM at 1-3 kV acceleration voltage and 8 mm working distance.

### Nanoindentation

To obtain the Young’s modulus of native tissues, PA hydrogels, and PA hydrogel-stabilized cardiac tissue slices pre- and post-DECIPHER, samples were attached to glass bottom petri dishes with superglue (for native tissue, GLUture was used instead, World Precision Instruments, 503763) and measured using the Optics11Life Chiaro Nanoindenter. To minimize sample-probe adhesion, 1% Pluronic (Sigma-Aldrich) solution was used to coat the probe prior to experiments. Matrix scans were performed on native tissue, PA-stabilized tissue, and DECIPHER samples with 500 μm step size (7×7) and 2 μm indentation depth. Matrix scans on plain PA hydrogel samples were performed with 800 μm step size (5×5) and 2 μm indentation depth. The Young’s modulus of each data point was obtained from the Hertzian contact model with Poisson’s ratio *ν* = 0.5 in Optics11 DataViewer software. n = 3-5 samples for each group.

### Primary cardiac fibroblast isolation

The use of animals was approved by Institutional Animal Care and Use Committee (IACUC), National University of Singapore. Mice were housed in individually ventilated cages, with sex-matched littermates, under standard conditions with a 12 hr light/dark cycle. Food and water were available ad-libitum. Primary young and aged cardiac fibroblasts were extracted from female young (1-2 months) and aged (24 months) C57BL/6J mice hearts, respectively, as previously described^19^. Briefly, mouse hearts were perfused with cardiac digestion buffers by intra-ventricular injection. Three rounds of gravity settling were applied to separate cardiomyocyte (CM) and non-CM cells. Non-CM cells were seeded onto untreated tissue culture plastic and washed after 1 hr to enrich for cardiac fibroblasts (CF). CFs were expanded in CF growth medium (DMEM/high glucose (Cytiva) supplemented with 15% fetal bovine serum (FBS, Gibco), Penicillin 100 U/mL, Streptomycin 100 µg/mL (Nacalai Tesque), FGF2 (10 ng/mL, Stemcell Technologies), SB-431542 (5 µM, Targetmol). All cells used in the *in vitro* sample reseeding studies were from passages 2 or 3. Prior to cell seeding, all DECIPHER samples were immersed in sterile 1X PBS overnight, UV sterilized in a biosafety cabinet (BSC) for 60 mins, and equilibrated in full maintenance media for 30 mins. The isolated cells were seeded at 10k cells/well in 1 mL of culture media (DMEM + 1% FBS + 1% pen/strep) onto DECIPHER samples of young or aged tissues stabilized in soft or stiff hydrogels and kept in a 37°C, 5% CO_2_, humidified incubator.

### Immunocytochemistry (ICC)

After culturing, cells were rinsed in sterile 1X PBS three times, then fixed with 4% formaldehyde for 15 mins, blocked with 2% BSA for 30 mins and permeabilized with 0.1% Triton X-100 for 15 mins for ICC. Samples were stained with primary antibodies (Table 2) overnight at 4°C in 2% BSA, rinsed three times in 1X PBS, followed by secondary antibodies and CF® Phalloidin for 1 hr at room temperature diluted in 2% BSA. The samples were washed with PBS three times (5 mins each) and stained with Hoechst 33342 in H_2_O for 10 mins at room temperature before three rounds of water rinses and mounted with Fluoromount-G^TM^.

### Low-input bulk RNA sequencing

All reagents mentioned are DNase/RNase/protease free and diluted with HyPure^TM^ Molecular Biology Grade Water (cytiva). Cells attached to the DECIPHER samples were lysed using TRIzol (Ambion) after scraping away the hydrogel region (to exclude any possible attached cells on the plain hydrogel region) and kept under −80°C pending RNA extraction. To further provide baseline (*in vivo*) control groups, freshly isolated young/aged primary CFs were directly lysed with TRIzol after extraction. Total RNA of samples was extracted using a chloroform-isopropanol approach according to the supplier’s instructions. Briefly, chloroform was added to lysed cell samples after thawing and vigorously mixed and centrifuged at 14,000 *×* g for 15 min before supernatant was collected. Cold isopropanol was then used to precipitate RNA, which was subsequently spun down at 14,000 *×* g for 15 min. Supernatant was discarded and RNA was washed twice with pre-cooled 75% ethanol followed by centrifugation at 14,000 *×* g for 10 min. The RNA samples were air dried for 5-10 mins before being dissolved in HyPure^TM^ water and processed for quality control with TapeStation High Sensitivity RNA ScreenTape Analysis. Sample RNA with an RNA integrity number (RIN^e^) larger than 8.5 (high quality) was applied to NEBNext^®^ Single Cell/Low Input RNA Library Prep Kit for Illumina^®^ for cDNA synthesis and sequence-ready library preparation according to supplier’s instructions. Specifically, 20 ng of total RNA from each sample was used for cDNA Synthesis and 8 PCR cycles were used for cDNA amplification. The libraries were indexed with NEBNext Multiplex Oligos for Illumina (96 Unique Dual Index Primer Pairs Set 2) and submitted to NovogeneAIT Genomics (Singapore) for sequencing on the Illumina NovaSeq 6000 system.

### Imaging and analysis

The IHC and ICC samples were observed using a Nikon A1R scanning laser confocal microscope (40x and 60x water immersion objectives) and subsequently imported to FIJI. To quantify ECM architectural changes post-decellularization, a FIJI macro, TWOMBLI, was used as previously described^20^. Three confocal images from pre- and post-decellularized samples were analyzed for collagen fiber alignment. Z-stack imaging was performed on Nile blue-tagged samples from the sample surface and reconstructed using Imaris. ICC samples stained with α-SMA and F-actin were analyzed with CellProfiler for intensity and morphological metrics. Nucleus/cytosol localization ratio of YAP was quantified using CellProfiler by identifying the fluorescence from a 30-pixel perinuclear ring as the cytosolic localization. Paxillin (focal adhesion) quantification was performed using a custom FIJI script.

### RNA-seq analysis

RNA-seq reads were aligned to mouse genome assembly mm39 with STAR (2.7.8a). The aligned reads were quantified using HTSeq (0.11.0) with Ensembl transcripts release 105. The above computation was performed within PartekFlow cloud environment with default parameters (v10.0.23.0425). Differential gene expression analysis was done with DESeq2 package^21^ (Bioconductor 3.17) and clusterProfiler package^22,23^ (Bioconductor 3.17) was used to perform functional enrichment analysis. Data visualization was performed using OriginLab 2023, GraphPad Prism 9, MATLAB R2023a, and TBTools^24^.

### Statistical analysis

Data are expressed as mean ± standard deviation (SD) unless otherwise specified. For transcriptomic analysis, standard error of the mean (SEM) was used. Two-way ANOVA or one-way ANOVA were applied for significance tests, where applicable, with multiple comparisons determined by preplanned Fisher’s least significant difference test addressing difference attributed to stiffness (SoftY vs. StiffY, SoftA vs. StiffA) or ECM age (SoftY vs. SoftA, StiffY vs. StiffA). Unless otherwise noted, all experiments were performed as independent triplicate experiments. Heatmaps demonstrating DEG analysis in RNA-seq data were graphed by averaging replicates with original (unaveraged) readings and dendrograms shown in Extended Data figures (except in Fig. 3b where original data is shown). Heatmap in Fig.3e is shown following the order of other DEG heatmaps, with the original clustering dendrogram shown in Extended Data Fig. 4a. Gene expression results for DECIPHER samples are normalized to the young and aged groups of their respective *in vivo* mimic, i.e., young CFs are normalized to young ECM in a soft matrix (YCFs on SoftY) while aged CFs are normalized to aged ECM in a stiff matrix (ACFs on StiffA).

## Results and Discussion

### DECIPHER maintains native ECM ligand presentation and enables independent stiffness tunability

The DECIPHER method (detailed in Fig. 1a) was inspired by a previously reported method for tissue clearing^16^, in which N-methylolacrylamide, formed by pre-reacting acrylamide hydrogel solution with formaldehyde, binds to the amine groups of tissue proteins. After crosslinking the polyacrylamide (PA) hydrogel with UV light, the tissue proteins are stabilized onto the PA mesh, thereby making an interconnected hybrid hydrogel, and then decellularized in place. We modified the original method in three ways: 1) by linking the PA gel to methacrylated coverslips, enabling stable sample handling; 2) by modifying the PA hydrogel solutions in order to dictate the scaffold stiffness; and 3) by optimizing the decellularization method to maintain ECM composition and architecture.

**Fig. 1.**
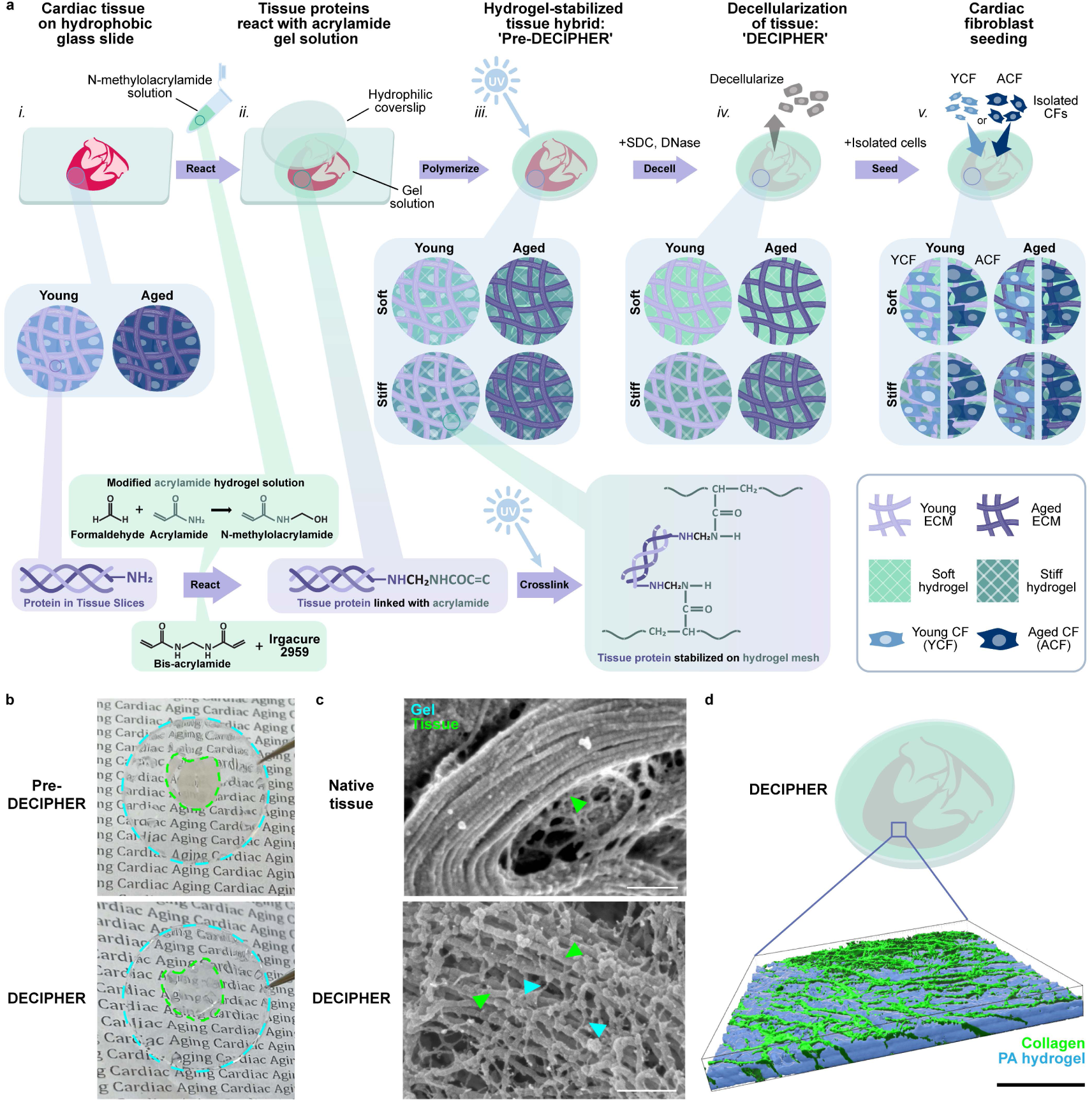
The DECIPHER system. Murine cardiac tissue (young and aged) was sliced with a vibratome, placed on a hydrophobic glass slide (*i.*), and incubated in hydrogel stabilization solution consisting of acrylamide pre-treated with formaldehyde, bis-acrylamide, and Irgacure 2959 of soft (~10 kPa) or stiff (~40 kPa) formulations (*ii.*). Tissue extracellular matrix (ECM) is indicated in purple (young – light purple, aged – dark purple) and hydrogel solutions in green (soft – light green, stiff – dark green). Crosslinking of the hydrogel mesh was carried out with UV (*iii*.). At this step, the tissue is stabilized in the hydrogel mesh (as indicated below in the green-to-purple box) and samples are referred to as ‘Pre-DECIPHER.’ Subsequent decellularization is carried out using SDC and DNase (*iv*.) and samples are referred to as ‘DECIPHER.’ The scaffolds are subsequently seeded with young or aged cardiac fibroblasts, YCFs (light blue) or ACFs (dark blue), respectively (*v*.) (**a**). Images of pre-DECIPHER and DECIPHER samples with coverslip/pure PA gel indicated by blue dashed lines and tissue scaffold indicated by green dashed lines (**b**). Scanning electron microscopy (SEM) images of native tissue and DECIPHER samples with green arrows indicating ECM fibers and blue arrows indicating the hydrogel mesh (**c**). 3D confocal reconstruction of DECIPHER samples with Nile Blue tagged acrylamide (blue) and labeled collagen (green) (**d**). Scale bars: 200 nm (**c**), 100 µm (**d**).

With DECIPHER, we created four sample combinations of young (1-2 mo) and aged (18-24 mo) ECM, stemming from the isolated native tissue, and young and aged stiffness, dictated by the hydrogel component. These samples are referred to as follows: ‘SoftY’ for soft, young ECM; ‘StiffY’ for stiff, young ECM; ‘SoftA’ for soft, aged ECM; and ‘StiffA’ for stiff, aged ECM (Fig.1 and Extended Data Fig. 1a). Decellularization increases transparency of the tissue and the DECIPHER sample is stabilized within the PA hydrogel on a coverslip (Fig. 1b). Scanning electron microscopy (SEM) of DECIPHER samples showed a crosslinked fibrous structure due to PA-stabilization of ECM fibers, which is not present surrounding collagen fibrils in native tissue (Fig. 1c). Furthermore, confocal imaging of Nile Blue-tagged acrylamide and CNA35 (collagen) reveals the penetration of PA into the tissue and surrounding ECM fibers leaving the ECM exposed on the surface for cell binding due to the placement on the hydrophobic glass slide during polymerization (Fig. 1d). This demonstrates that seeded cells will attach to the exposed ECM because unmodified PA hydrogels have been shown to repel cell adhesion^25^.

Immunohistochemistry (IHC) of nuclei and actin in young and aged tissues confirmed the complete removal of cellular structures while preserving ECM composition (collagen, fibronectin, hyaluronan, and laminin) (Fig. 2a, b). PicoGreen dsDNA assay (Extended Data Fig. 1c) further verified full decellularization^26^ while collagen and sGAG quantification showed minimal loss post-DECIPHER (>95.8% of collagen and >52.0% of sGAG were preserved; Extended Data Fig. 1d, e). The loss of sGAGs was ascribed to their removal on the plasma membrane and inside cells. After optimizing the decellularization approach to be minimally damaging using SDC and DNase, we confirmed collagen fibrils remained intact through staining with collagen hybridizing peptide (CHP) (Extended Data Fig. 1g). We compared our decellularization method to a widely-used decellularization protocol (SDS + Triton X-100) and found denatured collagen in both PA-stabilized and unstabilized samples (Extended Data Fig. 1g), although hydrogel stabilization was effective in reducing collagen denaturation. We next quantified architectural disruption using the TWOMBLI Fiji plugin^20^ of collagen IHC images and observed no signficant differences (Extended Data Fig. 1f, h).

**Fig. 2.**
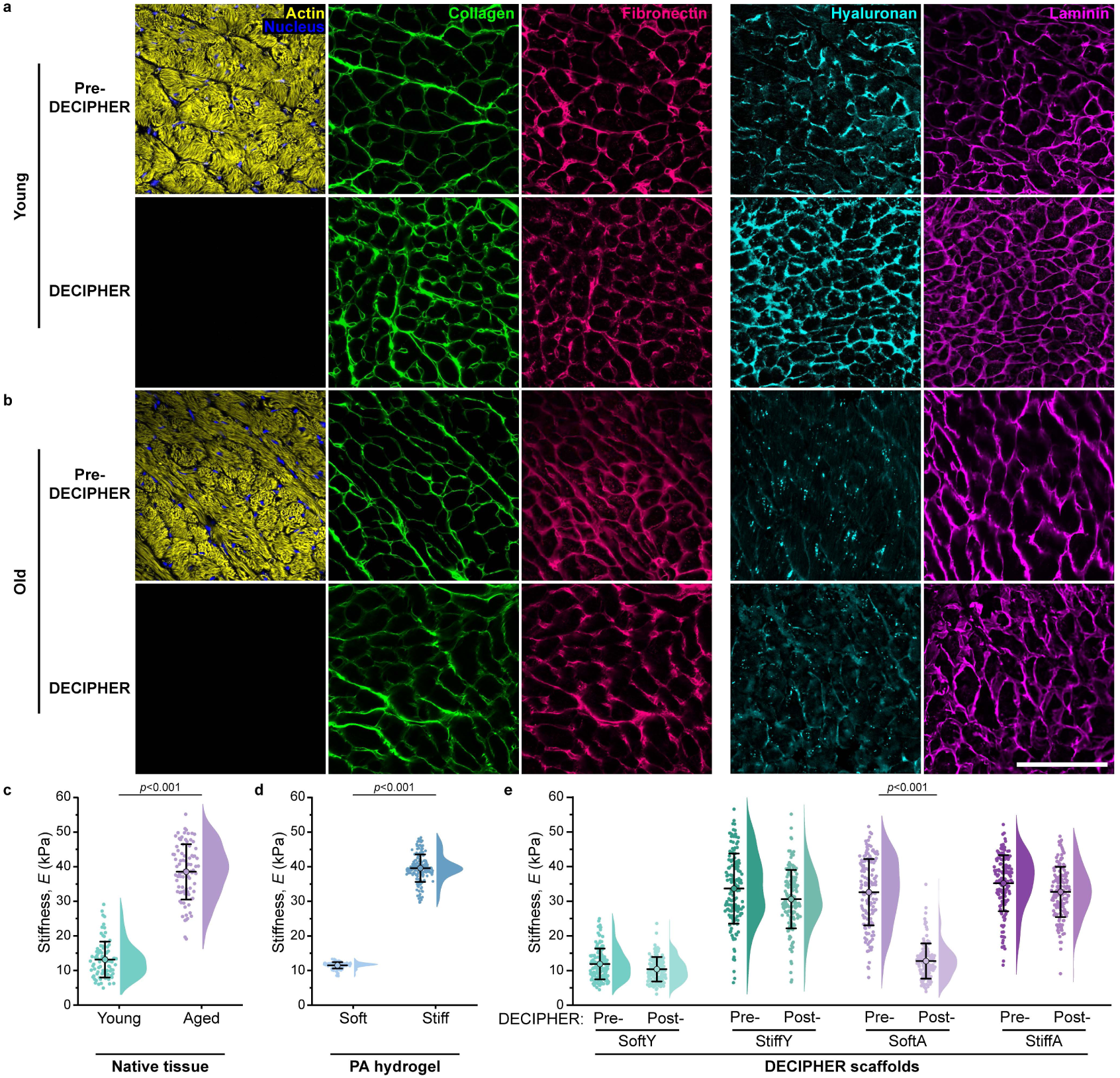
DECIPHER preserves ECM with tunable stiffness. IHC of pre-DECIPHER (top set) and DECIPHER (bottom set) samples from young (**a**) and aged (**b**) murine cardiac tissue for nucleus (blue), actin (yellow), collagen (green), fibronectin (cherry), hyaluronan (cyan), and laminin (magenta) show the removal of cellular structures in DECIPHER vs. pre-DECIPHER samples, as well as the preservation of ECM components. Nanoindentation of native cardiac tissue from young (teal) and aged (purple) mice (**c**) was used to dictate the PA hydrogel compositions for soft (light blue) and stiff (dark blue) samples (**d**). Stiffness mapping of DECIPHER samples showed mechanical tunability to maintain young stiffness in SoftY (teal), increase stiffness in StiffY (dark teal), decrease stiffness in SoftA (light purple, two-tailed t-test), and maintain stiffness in StiffA (dark purple) (**e**). Scale bar: 50 µm. n = 3 for (**c**, **e)** and n = 5 for (**d**).

One of the major improvements that DECIPHER brings to current decellularized ECM (dECM)-based material systems is *in situ* hydrogel-tissue stabilization. Hydrogel stabilization not only prevents tissue architecture disruption and damage during decellularization but also confers tunable mechanical properties to the dECM scaffold. Using nanoindentation, we first measured the stiffness (Young’s Modulus, *E*) of native tissue obtained from young or aged mice hearts (Fig. 2c). Young and aged tissue were found to be *E* = 13.1 ± 5.2 kPa and 38.6 ± 7.9 kPa, respectively, consistent with previous reports^1^. Subsequently, we optimized PA hydrogel compositions to mimic the measured tissue stiffnesses, *E* = 11.5 ± 0.9 kPa and 39.6 ± 4.0 kPa for young and aged tissue stiffness, respectively (Fig. 2d). Stiffness mapping of pre- and post-DECIPHER samples was carried out and demonstrated the ability to obtain decoupled stiffness tunability from native tissue stiffness (Fig. 2e, Extended Data Fig. 2). Specifically, young ECM can be tuned to aged stiffness and aged ECM can be tuned to young stiffness (Fig. 2e) due to the hydrogel component providing indentation resistance while dECM has been shown to exhibit reduced compression modulus and spontaneous shrinkage when unrestrained^27,28^. Taken together, DECIPHER allows for the mechanical tunability of decellularized tissue that is decoupled from native ECM composition and architecture.

### Global gene expression patterns in CFs are regulated by ECM as a function of age

Primary CFs were isolated from young (1 mo) and aged mice (24 mo) and lysed directly for RNA-seq. Isolated cells were examined for cell type-specific markers for Vimentin (fibroblast), CD31 (endothelial), and Troponin I (cardiomyocyte) confirming a pure fibroblast population in both age groups (Extended Data Fig. 3). Differential gene expression (DEG) analysis of aged CFs (ACFs) vs. young CFs (YCFs) found 490 upregulated genes and 597 downregulated genes (Fig. 3a), with heatmap representation showing differential expression patterns (Fig. 3b). Those upregulated included multiple senescence markers (e.g., Ccn1, Cdkn1a, and Cdkn2) and fibroblast-associated profibrotic markers (e.g., Ankrd1, Tnc, Ccr2, and Adamts1). The majority of significantly downregulated genes encode ECM proteins (e.g., Col1a1/2, Col3a1, Fn1, and Eln), which have been observed in aged rodent models^3^, as well as matricellular protein Ccn5, cyclin family genes (e.g., Ccnb1 and Ccnb2), and cyclin-dependent kinases (e.g., Cdk1 and Cdk6), indicating both the increased potential of ACFs to be activated^29^ and an abnormal proliferative ability that could suggest a senescent-like phenotype^30,31^ (Fig. 3a). KEGG enrichment analysis of the DEGs in ACFs vs. YCFs corroborated these findings as multiple significantly upregulated pathways indicated their greater potential toward profibrotic ECM remodeling, a senescent-like phenotype, and ECM deregulation as found in the aged heart^3^ (Fig. 3c).

**Fig. 3.**
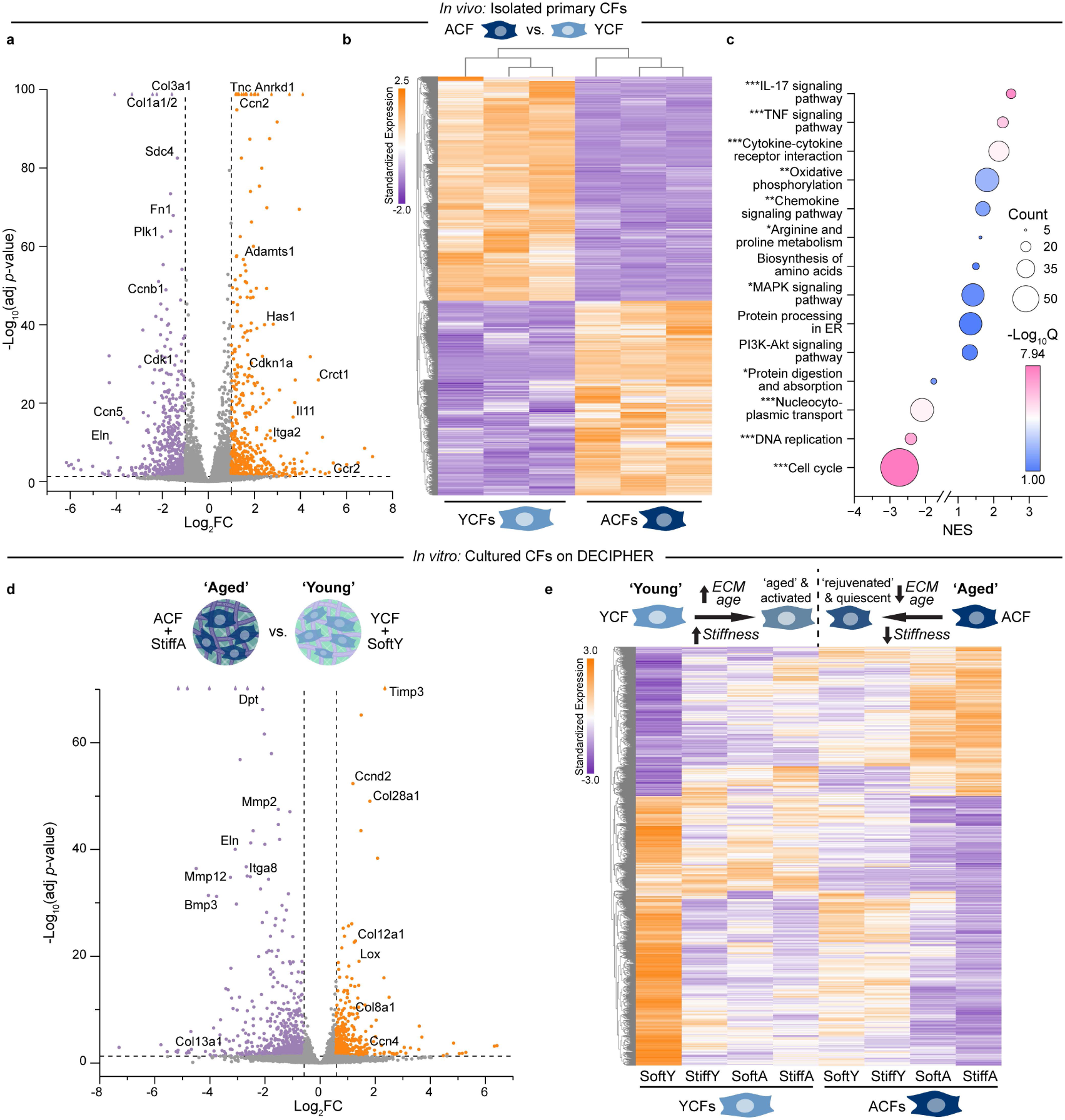
CFs exhibit age and ECM-dependent sensitivity. Differential gene expression (DEG) analysis of RNA-seq data of primary ACFs vs. YCFs show fold change (FC) expression with specific upregulated (orange) and downregulated (purple) genes highlighted (**a**). Heatmap of clustered DEGs with three sample replicates of YCFs and ACFs (**b**). KEGG enrichment analysis of ACF vs. YCF DEGs show activated and suppressed pathways. Bubble size indicates the number of genes differentially expressed in each pathway set and the color bar indicates the adjusted significance by Q-value. Significantly regulated pathways were marked with asterisks on the y-axis: *\*Q < 0.05*, *\*\*Q < 0.01*, *\*\*\*Q < 0.001*. (**c**). DEGs of CFs cultured on DECIPHER samples and compared as ACFs on StiffA (‘aged’ heart condition) vs. YCFs on SoftY (‘young’ heart condition) with upregulated (orange) and downregulated (purple) genes highlighted (**d**). Heatmap shows DEGs for YCFs and ACFs seeded on DECIPHER samples, which exhibit either an ‘aged/activated’-like shift for YCFs or ‘rejuvenated/quiescent’-like shift for ACFs as a function of stiffness and ECM (**e**). DEGs are FC >2 or < 0.5 and *p < 0.05* in (**a**) and (**b**); FC >1.5 or < 2/3 and *p < 0.05* in (**d**) and (**e**). Color bar in (**b**) and (**e**) indicate row-standardized expression.

Using our DECIPHER system, we next sought to understand how young and aged CFs perceive and respond to ECM stiffness and ligands, both individually and cooperatively. By seeding isolated CFs onto DECIPHER samples, we identified global DEGs (>1,100 genes) in the ‘aged’ state (i.e., ACFs cultured on a scaffold representing the aged heart, StiffA) vs. the ‘young’ state (i.e., YCFs cultured on a scaffold representing the young heart, SoftY) (Fig. 3d). Upregulated genes include those that encode matrix remodelers (e.g., Timp3 and Lox), specific ECM proteins (e.g., Col12a1 and Col8a1), and activators of fibroblasts (e.g., Ccn4 and Ccnd2), while downregulated genes include those that encode matrix degraders (e.g., MMPs 2 and 12), matrix adhesions (e.g., Dpt and Itga8), and specific ECM proteins (e.g., Eln and Col13a1).

When comparing ACFs on all other DECIPHER samples vs. StiffA, they exhibited a rejuvenation-like effect, shifting expression toward a quiescent phenotype, particularly on the young ECM samples (Fig. 3e, original clustering data in Extended Data Fig. 4a). The opposite trend was observed for YCFs on all other DECIPHER samples vs. SoftY, with expression indicating an aging-like shift and enhanced activation (Fig. 3e). Interestingly, ligand presentation had a larger impact vs. stiffness in driving young cells toward an aged state and aged cells toward a young state, with the greatest influence seen in aged cells.

We further applied comprehensive pairwise KEGG enrichment analyses on all DECIPHER samples to identify specific pathways that were activated or suppressed by stiffness cues as a function of age (Extended Data Fig. 5a) or by ECM as a function of age (Extended Data Fig. 5b). We found that ligand presentation promotes arginine and proline metabolism, while stiffness activates nucleocytoplasmic transport. In summary, primary ACFs exhibit both senescent and activated phenotypes, and young ECM is able to shift expression, regardless of stiffness, toward a rejuvenated-like state. Conversely, both higher stiffness and aged ECM can shift YCFs toward a more activated and aged-like state.

### Aged ECM ligand presentation outweighs aged mechanical cues in CF activation

The activation of CFs toward a myofibroblast phenotype was assessed after seeding YCFs and ACFs on DECIPHER samples with ICC for F-actin and *α*-smooth muscle actin (*α*-SMA). CFs generally exhibited an increase in cell area and decrease in circularity on stiffer substrates (Extended Data Fig. 6), while ECM age generally did not cause morphological changes. Furthermore, we observed enhanced expression of *α*-SMA and stress fiber (SF) formation as a function of stiffness regardless of ECM or cell age, albeit to differing levels (Fig. 4a, b, Extended Data Fig. 7a-d), corresponding to previous reports that stiffness alone can activate CFs^6,8,32^.

**Fig. 4.**
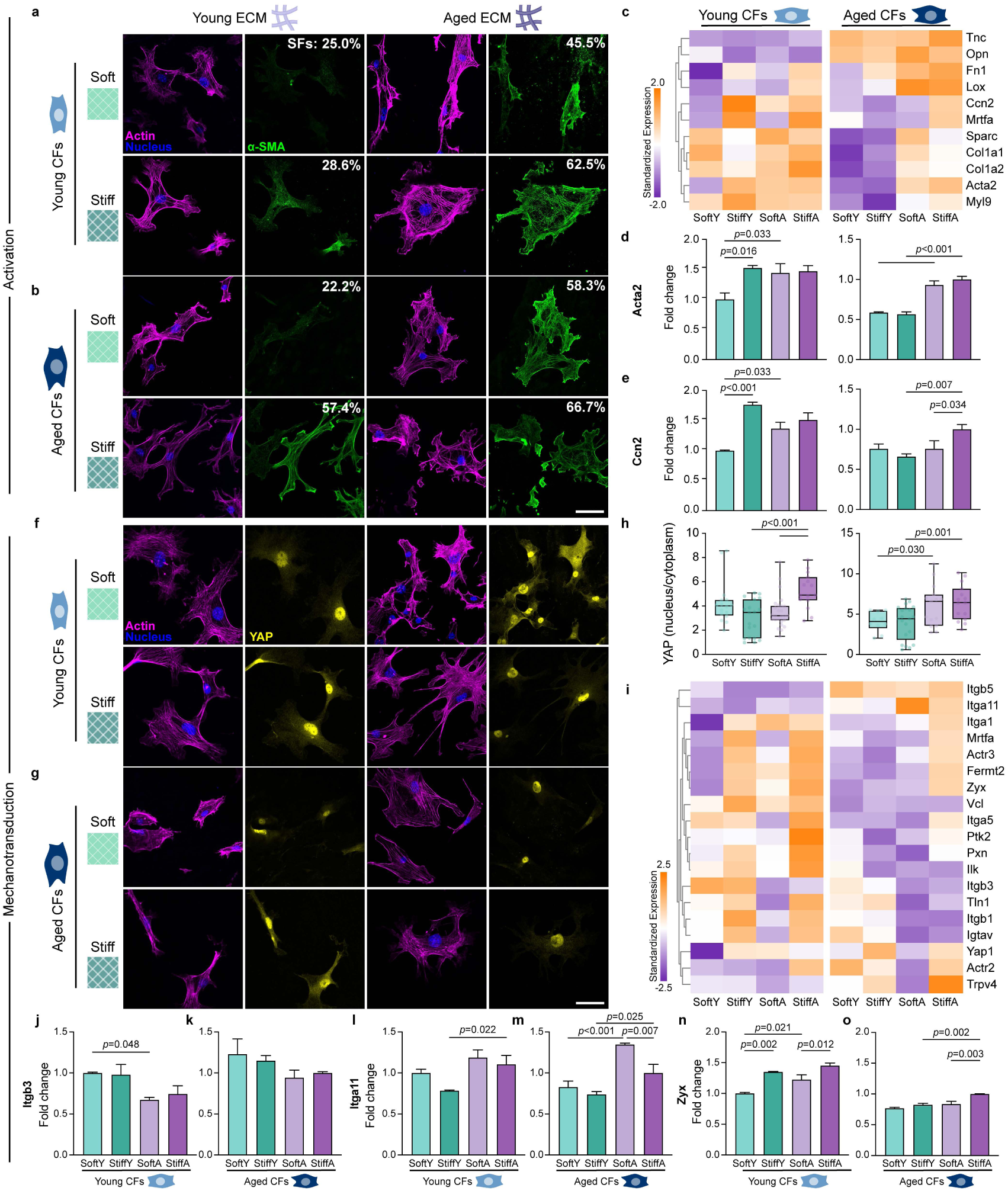
ECM ligand cues can override mechanical cues in CF activation and mechanotransduction. ICC of YCFs (**a**, **f**) and ACFs (**b**, **g**) on DECIPHER samples stained for nucleus (blue), F-actin (magenta), and *α*-SMA (green, in **a**, **b**) or YAP (yellow, in **f**, **g**). Activation analyses are shown in (**a**-**e**) and mechanotransduction analyses in (**f**-**o**). CF activation genes from RNA-seq data of YCFs and ACFs on DECIPHER samples with color bar indicating row-standardized expression (**c**). Normalized gene expression for CF activation markers (Acta2, Ccn2) for YCFs and ACFs on DECIPHER samples (**d**, **e**). YAP translocation (nuclear/cytoplasm) was quantified for YCFs and ACFs (**h**). RNA-seq analysis of CF mechanosensitive genes of YCFs and ACFs on DECIPHER samples with color bar indicating row-standardized expression (**i**). Normalized gene expression for highlighted integrins (Itgb3, 11) and focal adhesions (Zyx) in YCFs (**j**, **l**, **n**) and ACFs (**k**, **m**, **o**). Scale bars in (**a**, **b**, **f**, **g**) are 50 µm. DECIPHER samples in (**d**-**h** and **j**-**o**) are colored as follows: SoftY - light green, StiffY - dark green, SoftA - light purple, StiffA - dark purple.

Importantly, when comparing ECM ages, *α*-SMA was significantly reduced when CFs were plated on young ECM vs. aged ECM, indicating that the composition and architecture of young ECM can prevent CF activation, overriding the stiffness cue. This is particularly evident when YCFs are cultured on StiffY samples, as well as when ACFs are cultured on SoftY samples. However, for ACFs, young ECM alone is insufficient in preventing the upregulation of *α*-SMA as is seen on StiffY samples, highlighting the distinct mechanosensitive differences between young and aged cells. Aged ECM, on the other hand, enhanced *α*-SMA regardless of stiffness, with StiffA samples resulting in > 62% SF formation.

RNA-seq uncovered multiple genes that regulate CF activation in YCFs and ACFs as a function of ECM properties (Fig. 4c). Corresponding to the ICC data, both YCFs and ACFs upregulated Acta2 (encodes *α*-SMA) when on aged ECM and/or high stiffness except in the case of ACFs on StiffY (Fig. 4d). Similar correlations were observed for Ccn2 (encodes cellular communication network factor 2), which is another known CF activation marker (Fig. 4e). Myl9 (encodes myosin light chain 9), a downstream myofibroblast maturation marker, was significantly upregulated in ACFs on both SoftA and StiffA, pointing to the crucial role of ECM ligands in persistent CF activation independent of stiffness (Extended Data Fig. 7e, f). Taken together, we find that ECM ligand presentation can dictate myofibroblast activation state, outweighing the matrix stiffness cue.

### Aged CFs exhibit altered mechanosignaling

In addition to the well described cytokine-mediated route of CF activation and myofibroblast differentiation, the role of mechanical cues has recently been highlighted^6,31^. Thus, we investigated well-known mechanosensitive markers Yes-associated protein 1 (YAP) and paxillin with ICC on DECIPHER samples. Nuclear translocation of YAP, which is associated with fibroblast activation^33^, was enhanced on aged ECM (Fig. 4f-h). Although YAP translocation has been shown to depend on substrate stiffness, we found that young ECM supersedes mechanical cues in both young and ACFs, reducing the amount of nuclear translocation. This phenomenon could be attributed to the lower collagen I/III ratio in the young cardiac ECM that facilitates the degradation of translocated YAP^10^. Additionally, focal adhesion (FA) assembly was found to be limited on soft and/or young ECM, suggesting a less activated phenotype (Extended Data Fig. 8).

We next identified mechanosensitive targets from our RNA-seq data (Fig. 4i). YCFs activated focal adhesion proteins, serum response factor/myocardin-related transcription factor A (SRF/MRTF-A), and YAP on stiff and/or aged substrates (StiffY, SoftA, StiffA). ACFs, on the other hand, generally required both aged ECM and high stiffness (StiffA samples) to upregulate mechanosensitive proteins, which is in line with previously reported mechanosensing impairments seen in ACFs^10^. Despite altered mechanosignaling, we found that Actr2/3 was upregulated on StiffA samples, suggesting that ACFs favor facilitating ECM remodeling through actin-mediated force exertion^34^. Furthermore, the activation of transient receptor potential vanilloid 4 (TRPV4), which has previously been demonstrated to regulate the integration of multiple extracellular cues during activation and myofibroblast differentiation, was seen in ACFs on StiffA samples, suggesting their profibrotic potential^10,31^.

A wide spectrum of proteins regulating cell-ECM interactions exhibited varied expression to stiffness and young/aged ECM (Extended Data Fig. 9). The function of the arginine–glycine–aspartic acid (RGD)-binding β_3_ integrin in CFs has been previously identified in disease models to promote protein accumulation^31^, while its role in aging remains unclear. Here we found a decrease in β_3_ integrin expression in both young and aged CFs on aged ECM (Fig. 4j, k). We ascribe this to the overstretched fibronectin structure in the aged cardiac ECM that reduces available RGD sites^4^. Meanwhile, integrin α_11_ has been linked to CF activation and ECM protein synthesis in disease models^31^, and we observed a significant increase in α_11_ integrin expression among the cells on aged ECM (Fig. 4l, m). At the focal adhesion level, zyxin, which is responsible for actin regulation, was significantly upregulated in YCFs when exposed to either aged ECM and/or a stiff environment, while ACFs upregulated only when both cues were presented together (Fig. 4n, o). In summary, mechanotransduction in CFs is regulated by age state, with ACFs exhibiting altered mechanosignaling.

### Cell and matrix age determine ECM regulation

In the healthy heart, CFs maintain matrix homeostasis by balancing protein degradation, synthesis, and modification^31^. Thus, we sorted our RNA-seq data for ECM-regulating genes. ACFs on StiffA scaffolds showed an upregulation of matrix modifiers including lysyl oxidase (LOX) family enzymes and a downregulation of matrix degraders including matrix metalloproteases (MMPs), opposite from what is observed for YCFs on SoftY scaffolds (Fig. 5a-c, Extended Data Fig. 10). Enhanced LOX was also observed in ACFs on SoftA scaffolds, suggesting both the matrix and cell age increase the pro-fibrotic potential of CFs (Fig. 5a-c). Pro-fibrotic CFs increase crosslinking of ECM, which not only directly alters cellular behavior through architecture-mediated mechanisms^7,35^, but also exacerbates myocardial stiffening^3^. In addition, these samples showed significantly downregulated MMPs 2 and 3, which are responsible for normal tissue ECM turnover and degradation, respectively^36^ (Fig. 5a-c, Extended Data Fig. 10).

**Fig. 5.**
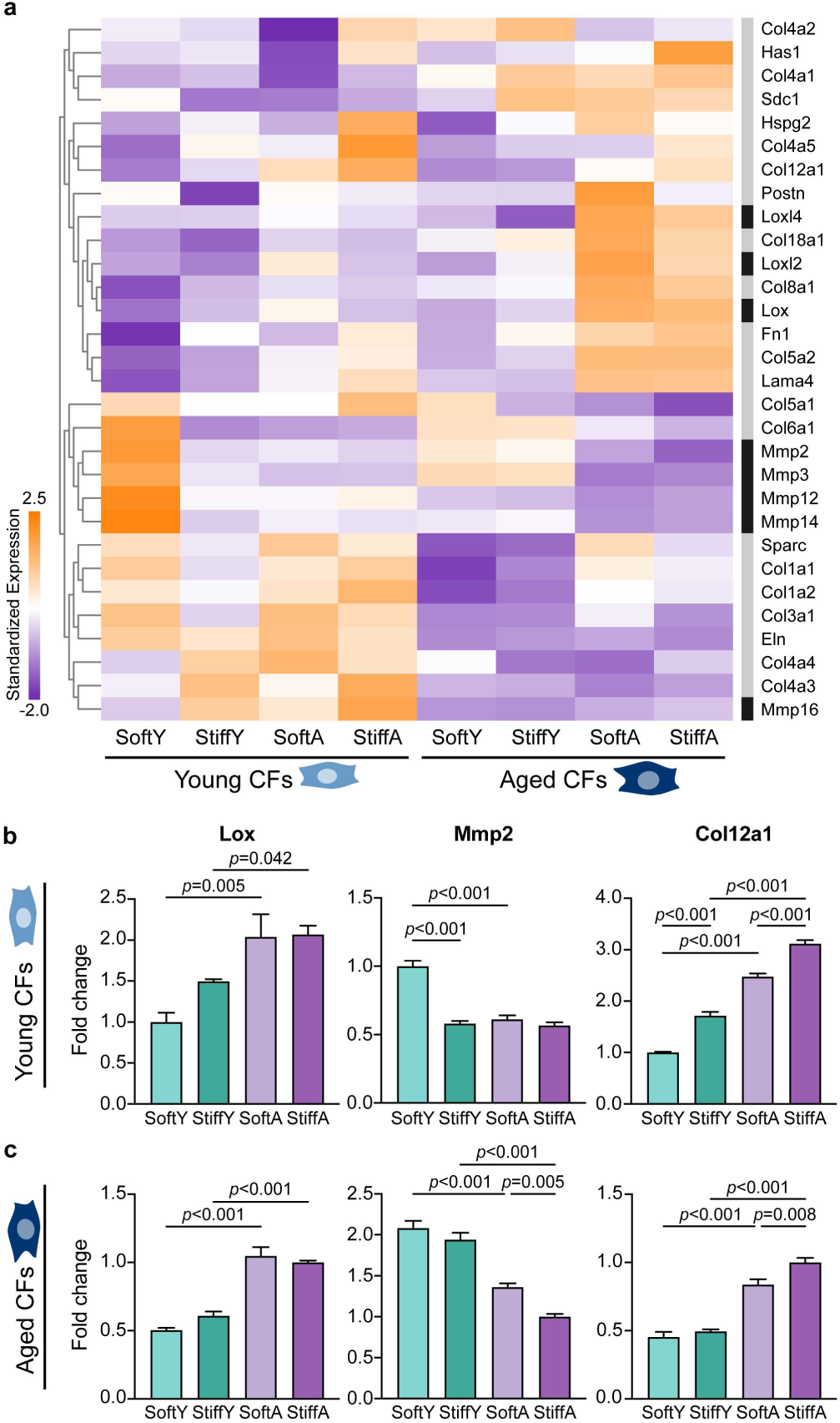
Regulators of ECM remodeling exhibit age-dependent expression. RNA-seq data of ECM degradation, crosslinking, and secretion genes of YCFs and ACFs on DECIPHER samples, with categorized genes indicated by the colored bars on the righthand side as grey: ECM protein secretion-related genes and black: matrix crosslinking or degradation-related genes. Color bar indicates row-standardized expression (**a**). Normalized gene expression for Lox, Mmp2, and Col12a1 for YCFs (**b**) and ACFs (**c**). DECIPHER samples in (**b, c**) are colored as follows: SoftY - light green, StiffY - dark green, SoftA - light purple, StiffA - dark purple.

The synthesis of multiple fibrillar and fibril-associated proteins (e.g., collagen I/III/V/XII, fibronectin) and non-fibrillar proteins (e.g., collagen VIII, laminin) were significantly upregulated in ACFs on SoftA and StiffA samples (Fig. 5a-c, Extended Data Fig. 11). However, collagen III expression decreased in a stiff environment, which could generate a positive feedback loop by increasing the collagen I/III ratio and consequently altering cardiac tissue mechanics^3^. The glycoprotein secreted protein acidic and rich in cysteine (SPARC), which is essential for ECM fibril formation and age-related tissue stiffening^3^, was also significantly upregulated in ACFs on SoftA and StiffA. Intriguingly, the synthesis of ECM proteins did not display a universal increase in response to aged ECM and high stiffness. Specifically, components associated with maintaining healthy mechanics and regulating fibrotic tissue size such as elastin^37^ and collagen V^5^ were downregulated (Fig. 5a). In summary, we find specific combinations of matrix mechanics and ECM age affect the extent of matrix regulation, yet CF and ECM age play the dominant roles.

### ECM ligands and mechanics cooperatively dictate CF senescence and rejuvenation

In the aging heart, senescence is an inevitable cellular phenotype that directly impacts organ physiology, leading to and/or accelerating cardiovascular diseases^30^. Due to the irreplaceable role of CFs in cardiac ECM deposition and remodeling, their acquisition of senescence is believed to cause ECM imbalance, especially during the wound healing process post-acute myocardial infarction (MI)^30^. In our global KEGG analysis (Extended Data Fig. 5), DECIPHER highlighted the roles of ECM and substrate stiffness in activating cellular senescence pathways, especially for the p53/p21 pathway which has been identified as one of the most well-characterized pathways in CF senescence^30,38^.

By comparing DEGS from DECIPHER groups seeded with young or aged CFs, we showed a shift of expression indicating *in vitro* ‘cardiac aging’ of YCFs from SoftY to StiffA, as well as ‘cardiac rejuvenation’ of ACFs from StiffA to SoftY (Fig. 6a, b). For YCFs, as the stiffness increases (SoftY to StiffY) or ECM ages (SoftY to SoftA), gene expression shifted toward a senescence-like phenotype that is seen on the StiffA substrate mimicking the aged heart (Fig. 6a, unaveraged heat maps in Extended Data Fig. 4b). For ACFs, young ECM and reduced modulus induced rejuvenation-like behaviors, with young ECM playing a greater role than reduced stiffness, as we observed for myofibroblast activation (Fig. 6b, unaveraged heat maps in Extended Data Fig. 4c). These findings reveal compelling prospects for rejuvenation therapies, particularly highlighting the potential of ECM-ligand targeted approaches.

**Fig. 6.**
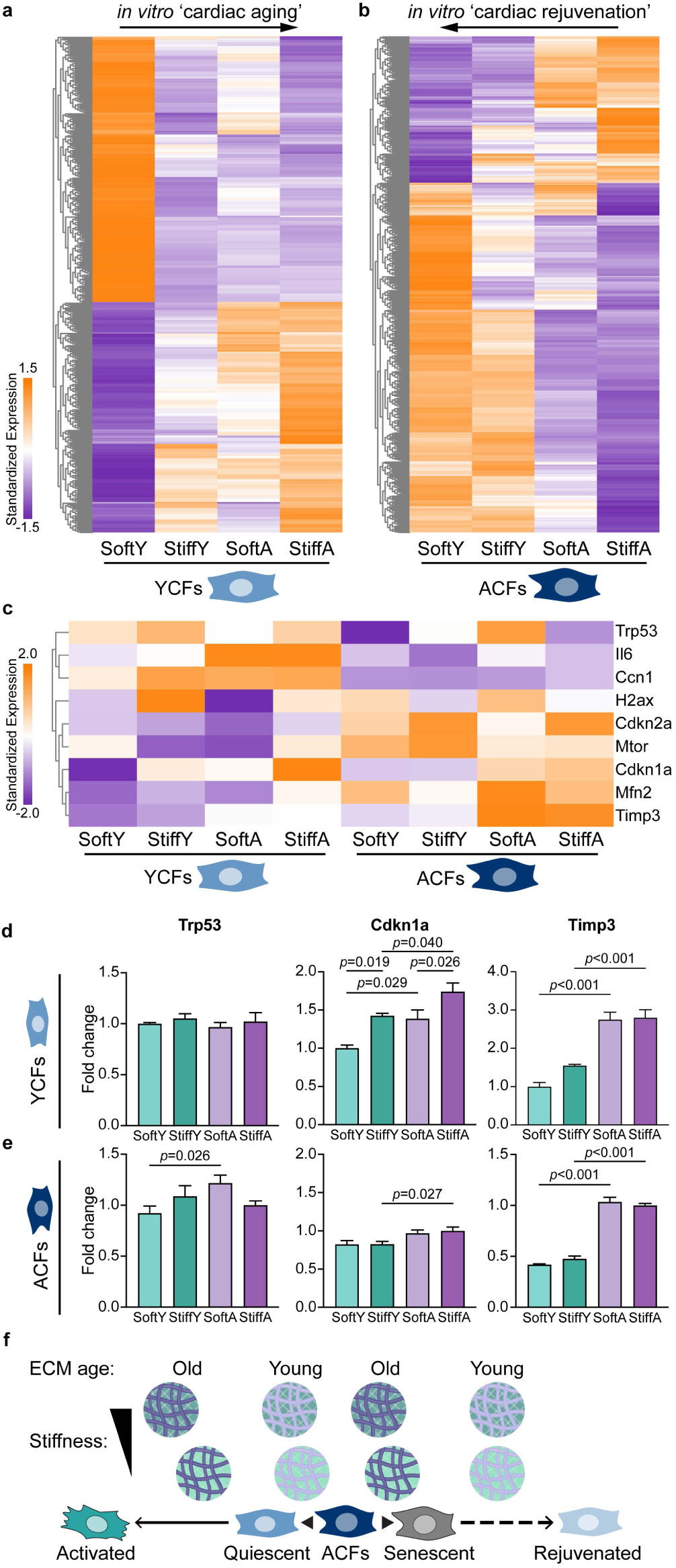
Cooperative ECM cues drive CF senescence and rejuvenation. Heatmaps of RNA-seq identified DEGs of YCFs (**a**) and ACFs (**b**) on StiffA vs. SoftY substrates with color bar indicating row-standardized expression. Heatmap of specific CF senescence regulators with color bar indicating row-standardized expression (**c**). Normalized gene expression for Trp53, Cdkn1a, and Timp3 for YCFs (**d**) and ACFs (**e**). Depiction of ECM property regulation (ECM ligand age vs. stiffness) of ACF activation, senescence, and rejuvenation as elucidated via DECIPHER (**f**). DEGs are FC >1.5 or < 2/3 and *p < 0.05* in (**a**) and (**b**). DECIPHER samples in (**d**, **e**) are colored as follows: SoftY - light green, StiffY - dark green, SoftA - light purple, StiffA - dark purple.

Although CFs on StiffA DECIPHER samples did not demonstrate a terminally senescent phenotype regarding collagen expression (reduced) and proinflammatory cytokines expression (enhanced)^30,39^, many key senescence-associated pathways were activated including p53/p21 (encoded by Trp53/Cdkn1a), p16 (encoded by Cdkn2a), and p38 mitogen-activated protein kinase (MAPK) (Fig. 6c-e). Downstream expression of senescence-associated secretory phenotype (SASP) proteins that have been shown in senescent fibroblasts^30,40^, including tissue inhibitor of metalloproteinase 3 (TIMP-3), cellular communication network factor 1 (CCN1) and interleukin 6 (IL-6), were also significantly enhanced (Fig. 6c-e, Extended Data Fig. 12). These results suggest the critical roles of aged ECM and myocardial stiffening in inducing CF senescence, both individually and cooperatively (Fig. 6f).

## Outlook

Important studies have recently uncovered the essential roles of ECM composition, architecture, and mechanics in driving cellular function^2,6,35^. To describe the impact of ECM-specific properties, new tunable dECM-based biomaterials systems have been designed for a variety of applications including 2D and 3D *in vitro* bulk hydrogel systems, injectable *in vivo* hydrogel formulations, and complex bioprinted scaffolds^2,41–43^. While dECM-based biomaterials can provide faithful recapitulation of native ECM composition to some degree, preserving mechanical and architectural aspects of the native tissue remain a challenge. This is crucial because ECM architecture has been shown to regulate cellular phenotypes and organ physiology in recent studies^7,35,44,45^, while ECM mechanics have become increasingly appreciated in the field of cell biology over the past two decades^46–48^. Although some dECM-based biomaterial systems can provide tunable stiffness, albeit within a limited range and typically much softer than the original tissue stiffness, the tunability depends on other ECM properties. For instance, when modifying mechanics of lyophilized dECM, ECM structure and composition will be influenced. Thus, our new approach (DECIPHER) not only faithfully mimics the native ECM composition and architecture, but also provides independent mechanical tunability spanning a greater range of physiologically relevant stiffnesses due to the fact that the hydrogel component can be synthesized from ~1 to 100 kPa.

Here, we show that we can decouple the contributions of age-specific ECM ligand presentation and age-specific stiffness on cell behavior (Fig. 1). Importantly, we find that biochemical signaling and ligand presentation from the ECM can override stiffness-dependent mechanosignaling in age-related CF activation, matrix remodeling, and acquisition of senescent-like phenotypes (Fig. 6f), highlighting the key roles of ligand-specific cues that have been shown to dictate other cellular behaviors, including collective migration and stem cell fate^49,50^. More remarkably, we show that young ECM contributes to the rejuvenation potential of ACFs through distinct signaling pathways. The ultimate application of this platform is to identify new targets for matrix-based rejuvenation strategies, and we found that young, soft ECM could enhance the rejuvenation potential of ACFs. How CFs integrate biochemical, architectural, and mechanobiological signaling to determine their phenotypical transition toward an aged, dysfunctional state still requires further investigation with this system, and future studies will focus on the dynamics of this process, as previous reports have highlighted the importance of time dependence in cardiac function^51,52^. Taken together, we describe here a novel material platform that has widespread applicability to other tissues and/or disease types for which cell-ECM interactions play a crucial role.

## Acknowledgements

The authors thank Jennifer Marlena (MBI, NUS) for illustrations; Jingjing Zheng (Dept. of Chemistry, Southern University of Science and Technology - SUSTech), Hepi Hari Susapto, Xu Gao, Jashan Preet Singh, and Ranmadusha Merengha Hengst (MBI, NUS) for fruitful discussions and technical advice; Chii Jou Chan, Vinod S/O Prabhakaran, and Ng Boon Heng (MBI, NUS) for providing HABP and vibratome for usage; and Ong Hui Ting (MBI, NUS) for image analysis codes. We thank the MBI Wet Lab Core for facility and technical support, the Singapore Microscopy and Bioimaging Analysis (SiMBA, MBI) core for microscopy and data processing facilities, and the High-throughput Molecular Genetics (HMG, MBI) core for RNA-seq support.

This work was supported by the Ministry of Education under the Research Centres of Excellence programme through the Mechanobiology Institute at the National University of Singapore and the Biomedical Engineering Department at the National University of Singapore, as well as the Singapore Ministry of Education Academic Research Fund Tier 3 (MOE Grant No: MOET32021-0003) to J.L.Y.

## Author contributions

J.L.Y. and A.R.S. conceptualized the project and wrote the manuscript. J.L.Y. supervised data collection and analysis. A.R.S. performed the majority of experiments and data analysis. M.F.H.R., X.S., and D.C. assisted in bench experiments. M.F.H.R. and D.C. assisted in data analysis. M.A-J. performed primary cell isolations. J.Z. assisted in RNA-seq experiments and analysis. R.S.F. provided key reagents and contributed toward discussion.

## Data availability

The data supporting the findings of this study are available within the article and supplementary materials. RNA-seq raw data will be uploaded to NCBI GEO database prior to publication with accession code provided.

## Code availability

All codes and scripts used in this study are available from the corresponding author on request.

## Supplemental figures

Dummy

**Extended Data Fig. 1.**
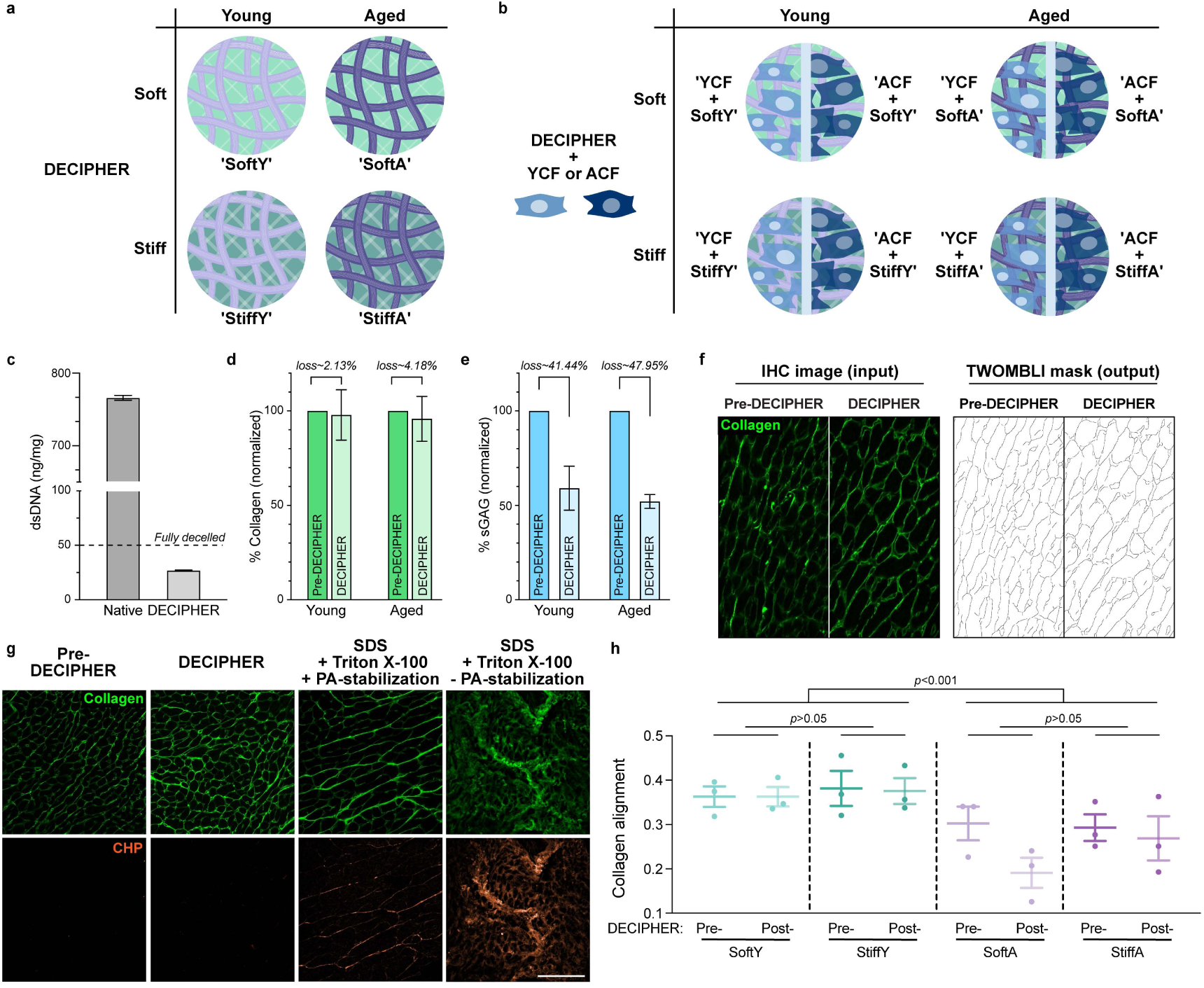
Characterization of DECIPHER samples. Nomenclatures used in this study describing DECIPHER samples before (**a**) and after cell seeding (**b**). Assay for dsDNA content in native tissue (‘Native’) vs. DECIPHER samples, with the accepted decellularization threshold of 50 ng/mg indicated by the dashed line (**c**). Hydroxyproline (collagen) content (**d**), and sGAG content (**e**) in Pre-DECIPHER (green in **d**, blue in **e**) vs. DECIPHER (light green in **d**, light blue in **e**) samples. Representative images for TWOMBLI analysis with IHC images stained for collagen (green) pre- and post-DECIPHER (input) and the resulting fiber maps (output) (**f**). IHC images of collagen hybridizing peptide (CHP, orange) and collagen (green) stained samples of Pre-DECIPHER, DECIPHER, SDS + Triton X-100 +/- PA stabilization (**g**). Collagen alignment indicated in pre- and post-DECIPHER samples for SoftY (light teal), StiffY (teal), SoftA (light purple), and StiffA (purple) from TWOMBLI analysis of images in (**f**) followed by two-tailed Student’s t-test (**h**). Scale bar is 50 µm.

**Extended Data Fig. 2.**
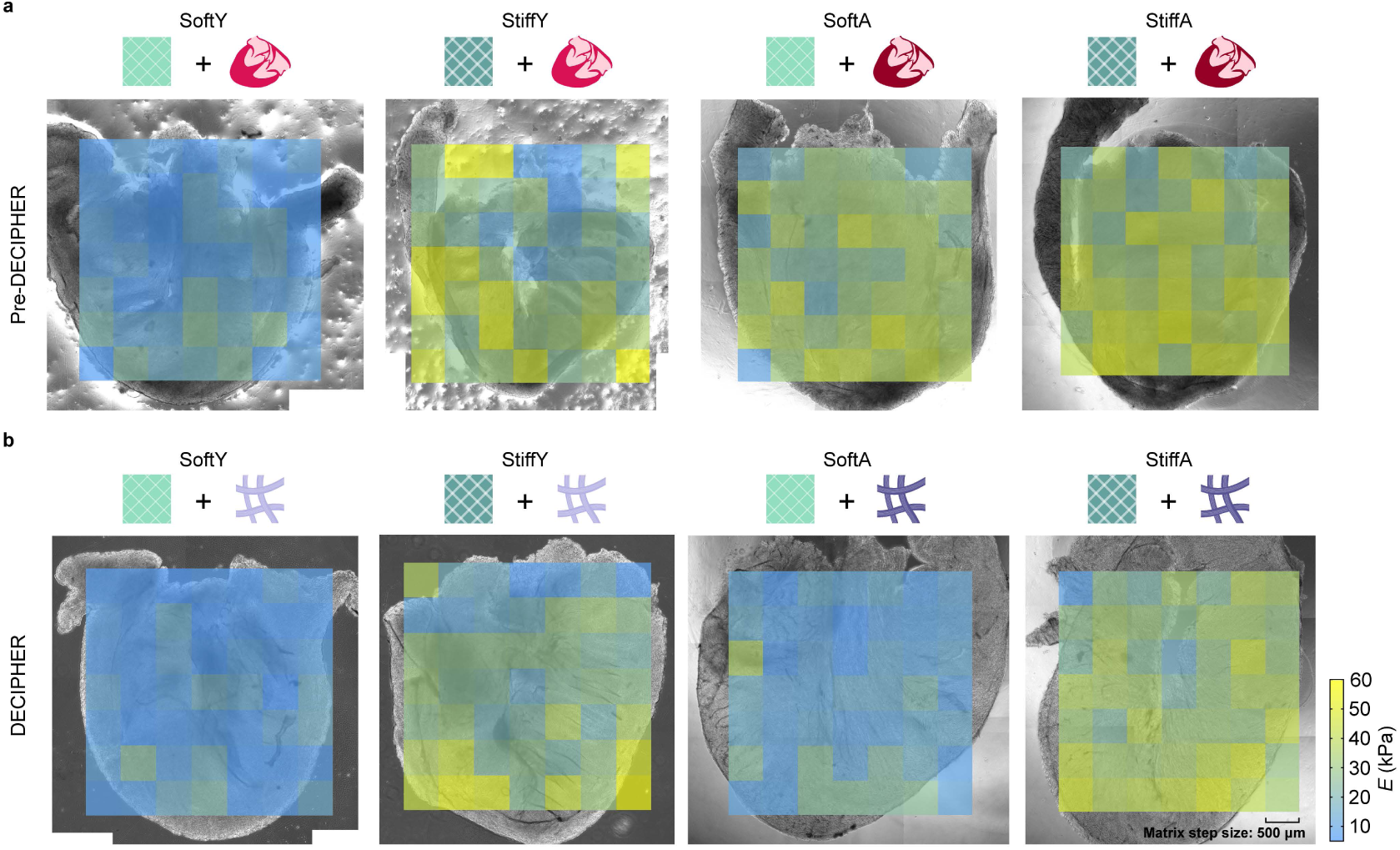
Stiffness mapping of pre- and post-DECIPHER samples reveals scaffold tunability. Nanoindentation maps of Pre-DECIPHER samples of young (Y) and aged (A) tissue stabilized in soft and stiff PA hydrogels (**a**) and of DECIPHER samples (**b**). Icons for PA gel stiffness are: light green box for Soft and dark green box for Stiff. Icons for heart age are: red heart for Young and dark red heart for Aged (Pre-DECIPHER). Icons for ECM age are: light purple for Young and dark purple for aged (DECIPHER). Color bar indicates the Young’s modulus, *E* ranging from 5 kPa (blue) to 60 kPa (yellow). Step size of measurements is 500 µm, indicated by the small squares in the maps.

**Extended Data Fig. 3.**
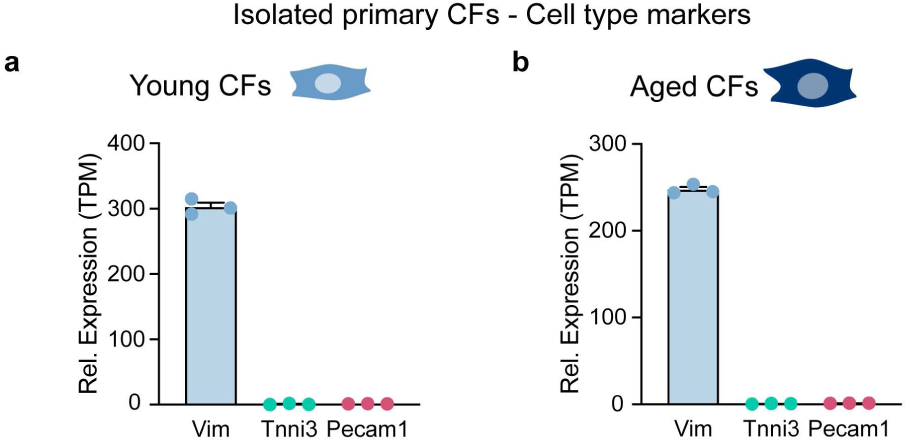
Specificity of primary CF isolation. RNA-seq reveals expression levels of Vimentin (Vim, blue), Troponin-I (Tnni3, green), and CD31 (Pecam1, red). A pure Vim-expressing fibroblast population was demonstrated with negligible Pecam1-expressing endothelial cells and Tnni3-expressing cardiomyocytes in both young (**a**) and aged (**b**) isolated CFs.

**Extended Data Fig. 4.**
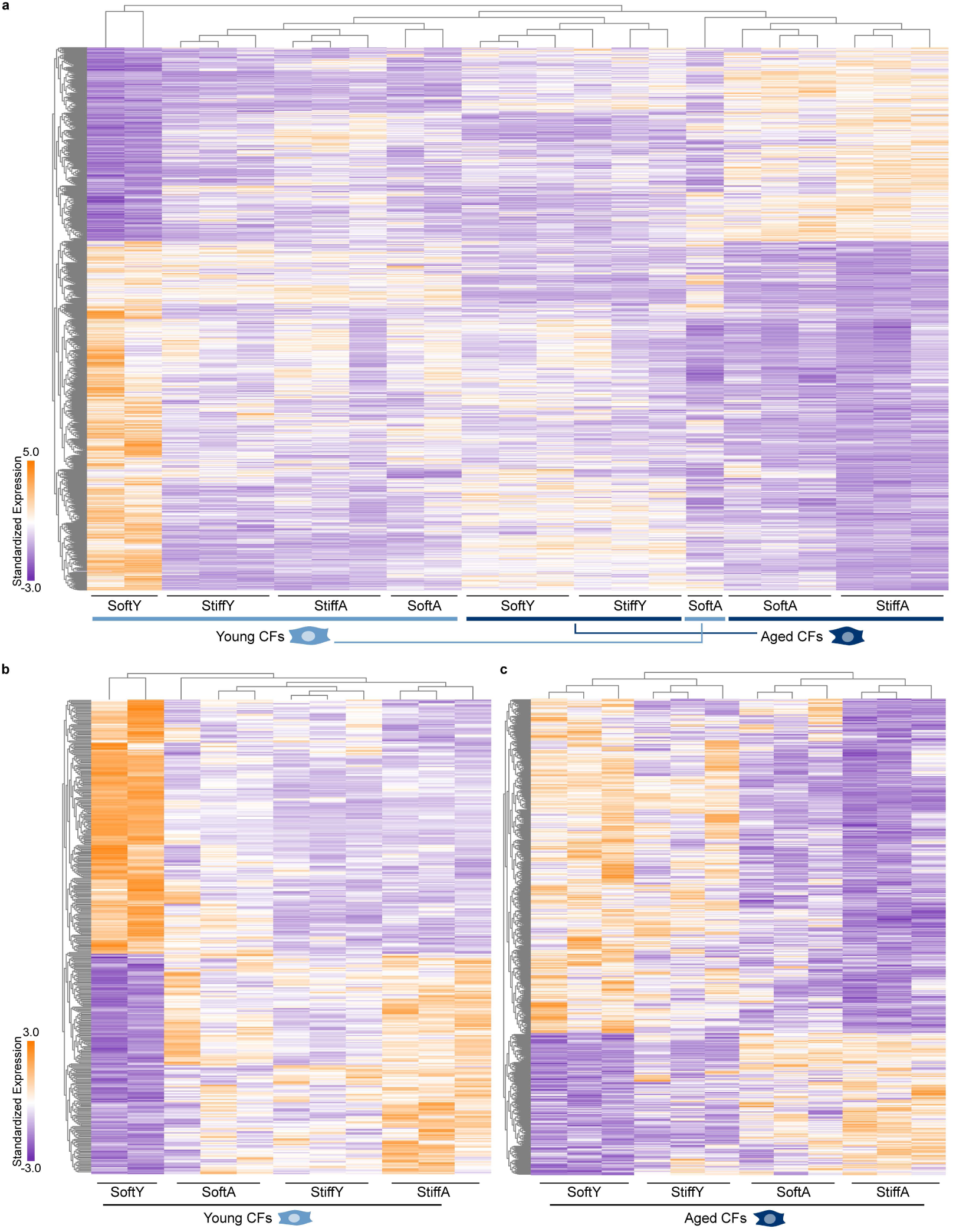
CF activation/quiescence and aging/rejuvenation-like phenotypes shown by original DEG heatmaps with dendrograms. Sample replicates are non-averaged in this graph set, where **a**, **b**, **c** correspond to Fig. 3d, Fig. 6a, and Fig. 6b, respectively. DEGs are FC >1.5 or < 2/3 and *p < 0.05*.

**Extended Data Fig. 5.**
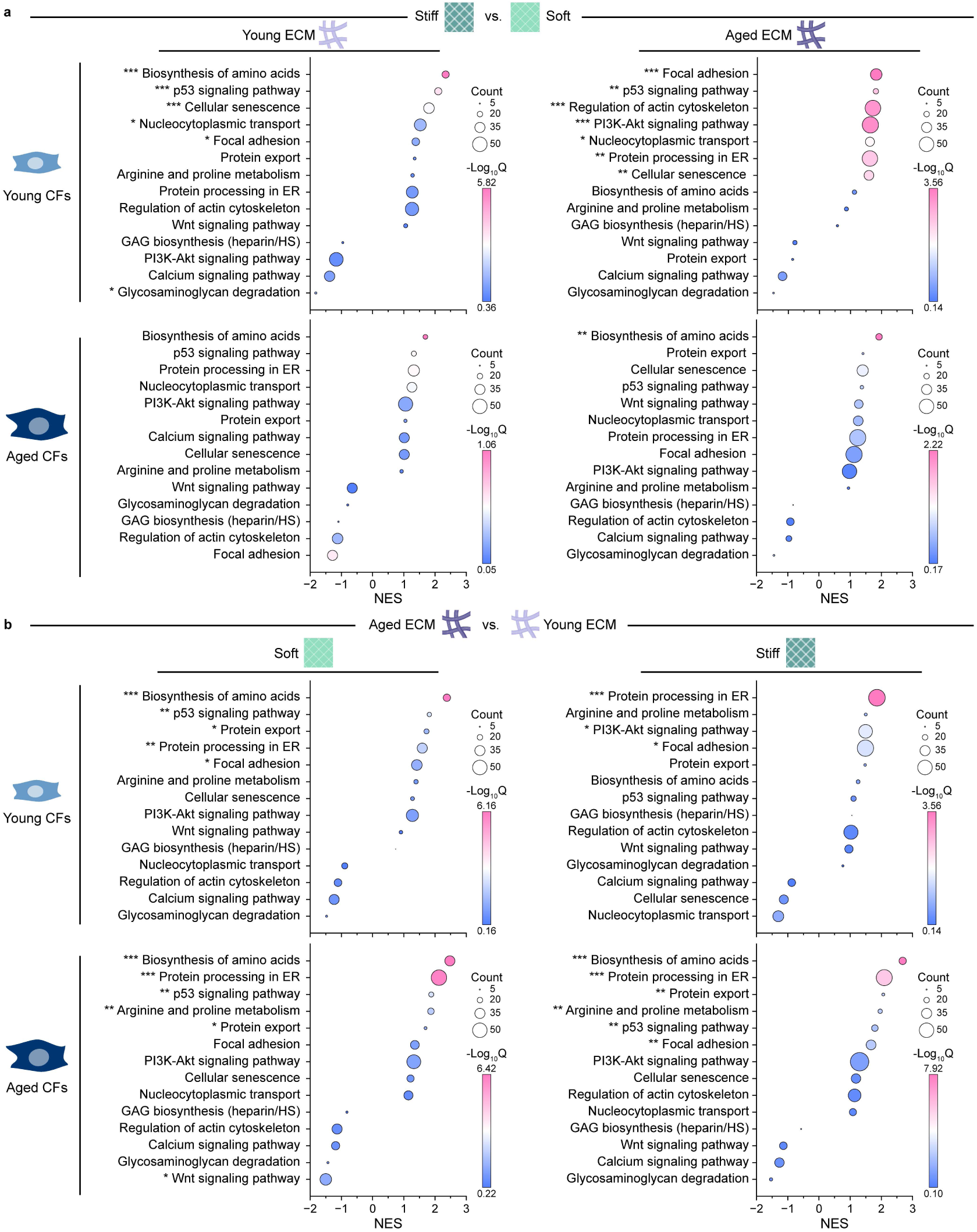
KEGG enrichment analysis of CFs seeded on DECIPHER samples. Highlighted pathways of interest in young or aged CFs impacted by mechanical cues (Stiff vs. Soft, **a**) or ECM aging (Aged ECM vs. Young ECM, **b**) were graphed in the order of normalized enrichment score (NES) with color bar showing adjusted significance using Q-value. Significantly regulated pathways were marked with asterisks beside the pathway name: *\*Q < 0.05*, *\*\*Q < 0.01*, *\*\*\*Q < 0.001*. Bubble size indicates the number of genes differentially expressed in each pathway set.

**Extended Data Fig. 6.**
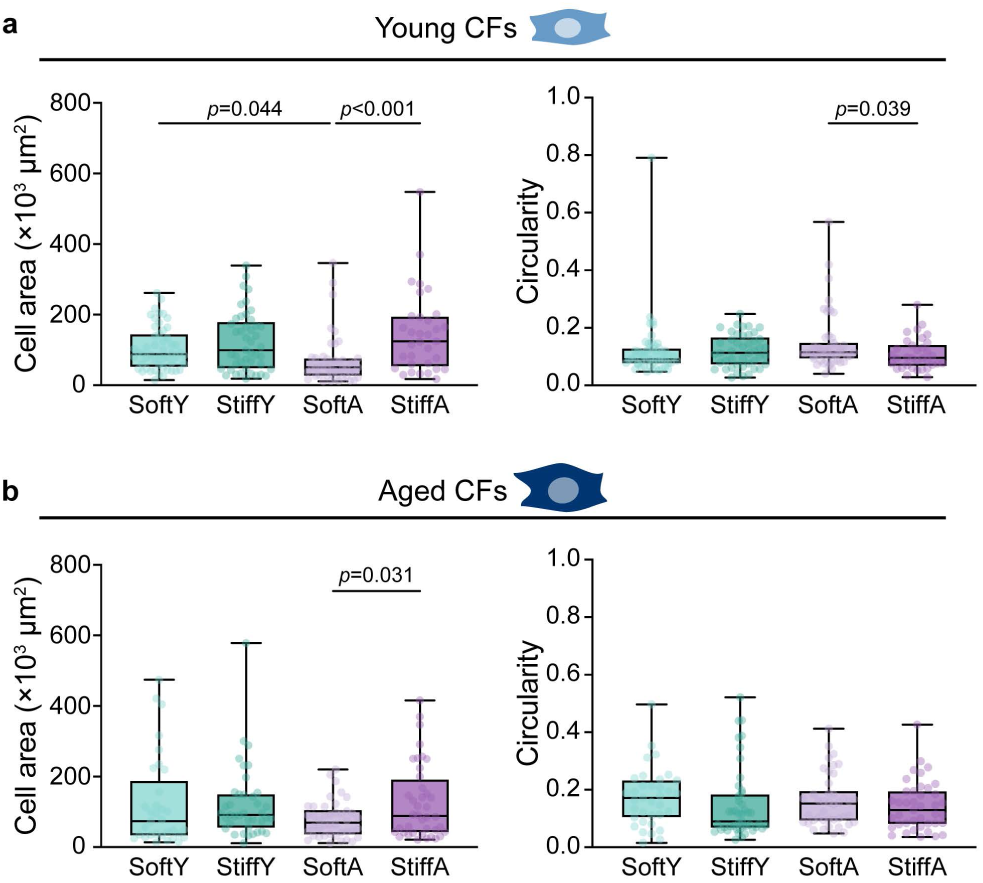
Morphological changes in CF activation are governed by stiffness. Cell area and circularity of YCFs (**a**) and ACFs (**b**) on DECIPHER samples quantified by F-actin ICC images. Cell data on DECIPHER samples are colored as follows: SoftY - light green, StiffY - dark green, SoftA - light purple, StiffA - dark purple.

**Extended Data Fig. 7.**
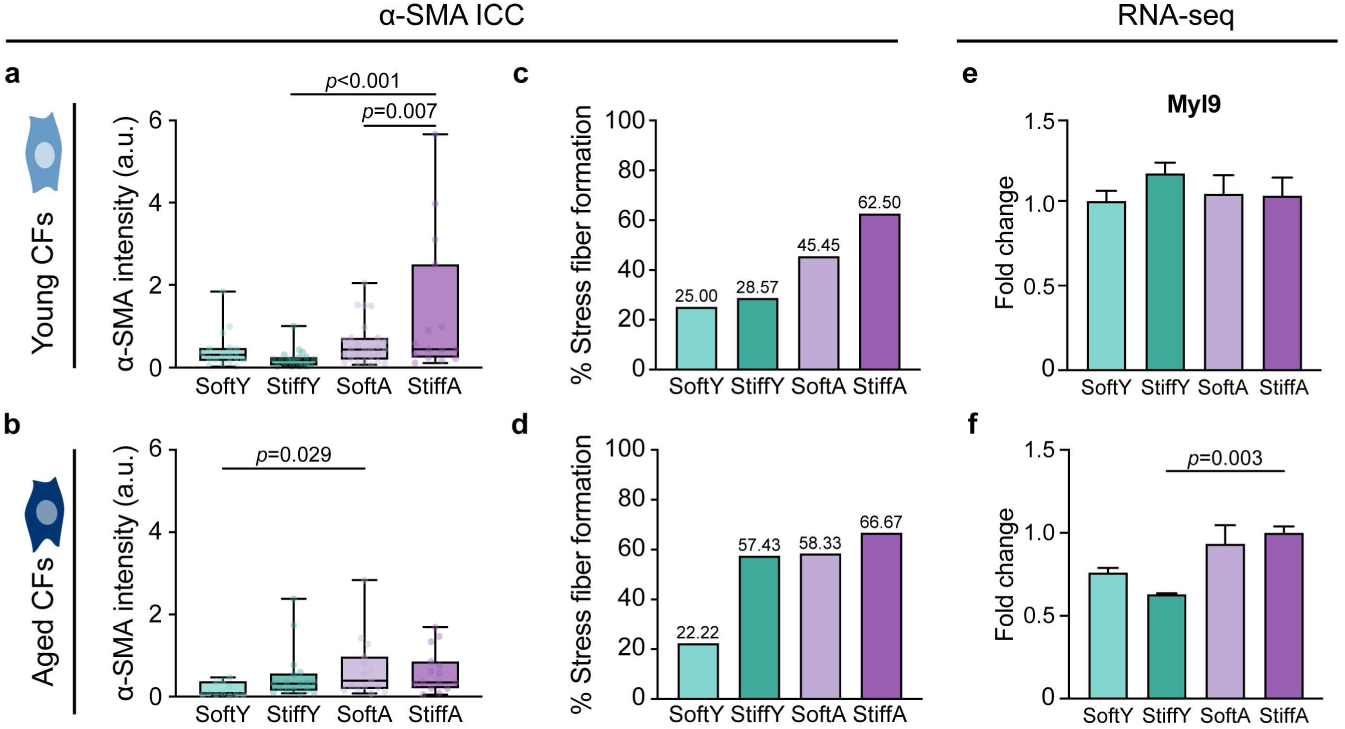
ECM ligands outweigh stiffness in modulating key CF activation phenotypes and gene expression. ICC images (shown in Fig. 4) quantified for *α*-SMA intensity and stress fiber assembly for YCF (**a**, **c**) and ACF (**b**, **d**). Expression of additional end-stage myofibroblast maturation marker MYL9 in YCFs (**e**) and ACFs (**f**). Cell data obtained from different DECIPHER samples are colored as follows: SoftY - light green, StiffY - dark green, SoftA - light purple, StiffA - dark purple.

**Extended Data Fig. 8.**
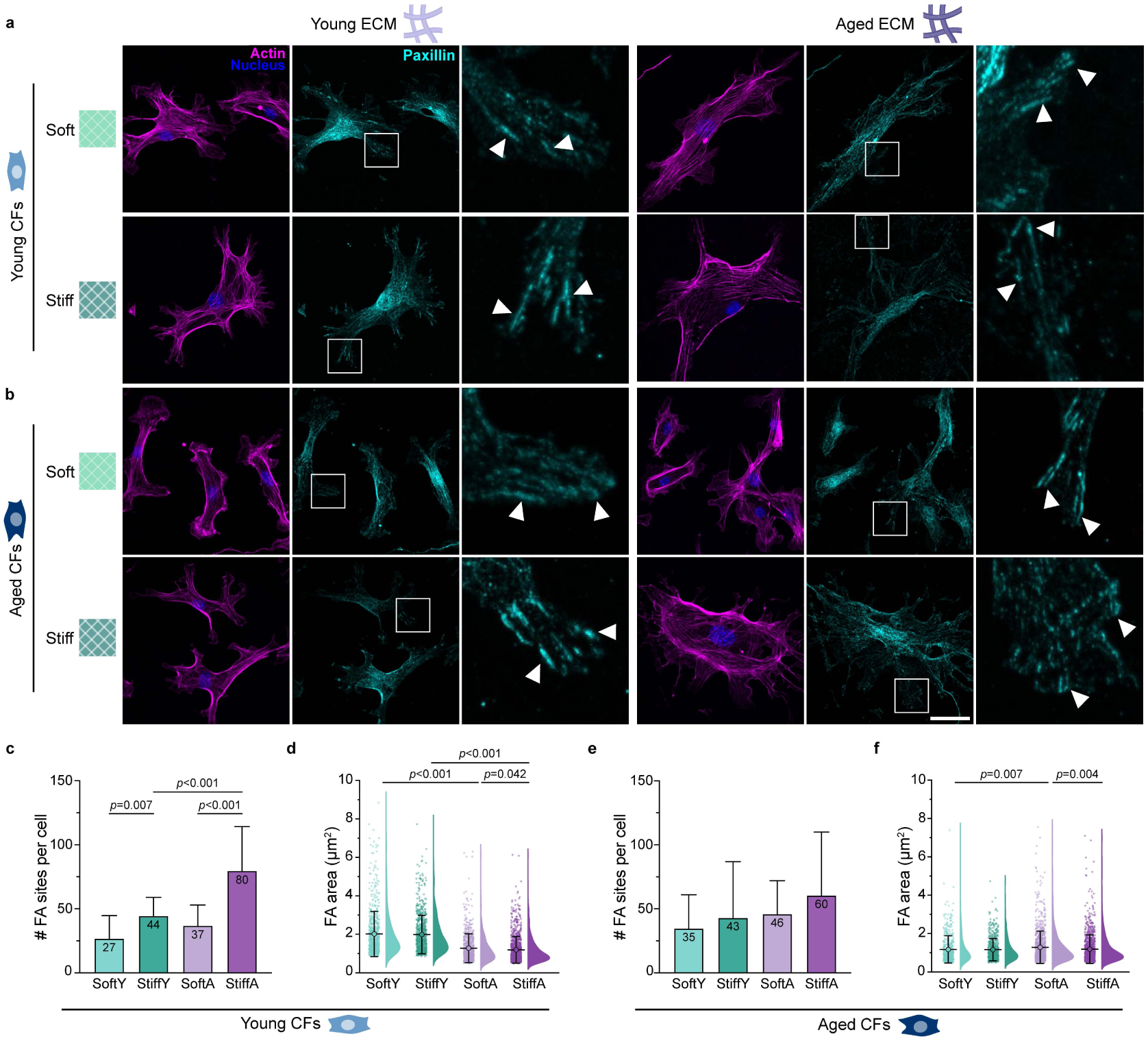
Focal adhesion formation on DECIPHER scaffolds. ICC of F-actin (magenta), nucleus (blue), and paxillin (teal) in YCFs (**a**) and ACFs (**b**) on DECIPHER samples. Paxillin insets are indicated by the white boxes, and white arrows point to focal adhesions. The number and area of focal adhesion sites in YCFs (**c, d**) and ACFs (**e, f**) quantified from paxillin staining. ICC data on different DECIPHER samples in (**c**, **d**, **e**, **f**) are colored as follows: SoftY - light green, StiffY - dark green, SoftA - light purple, StiffA - dark purple. Scale bar: 50 µm.

**Extended Data Fig. 9.**
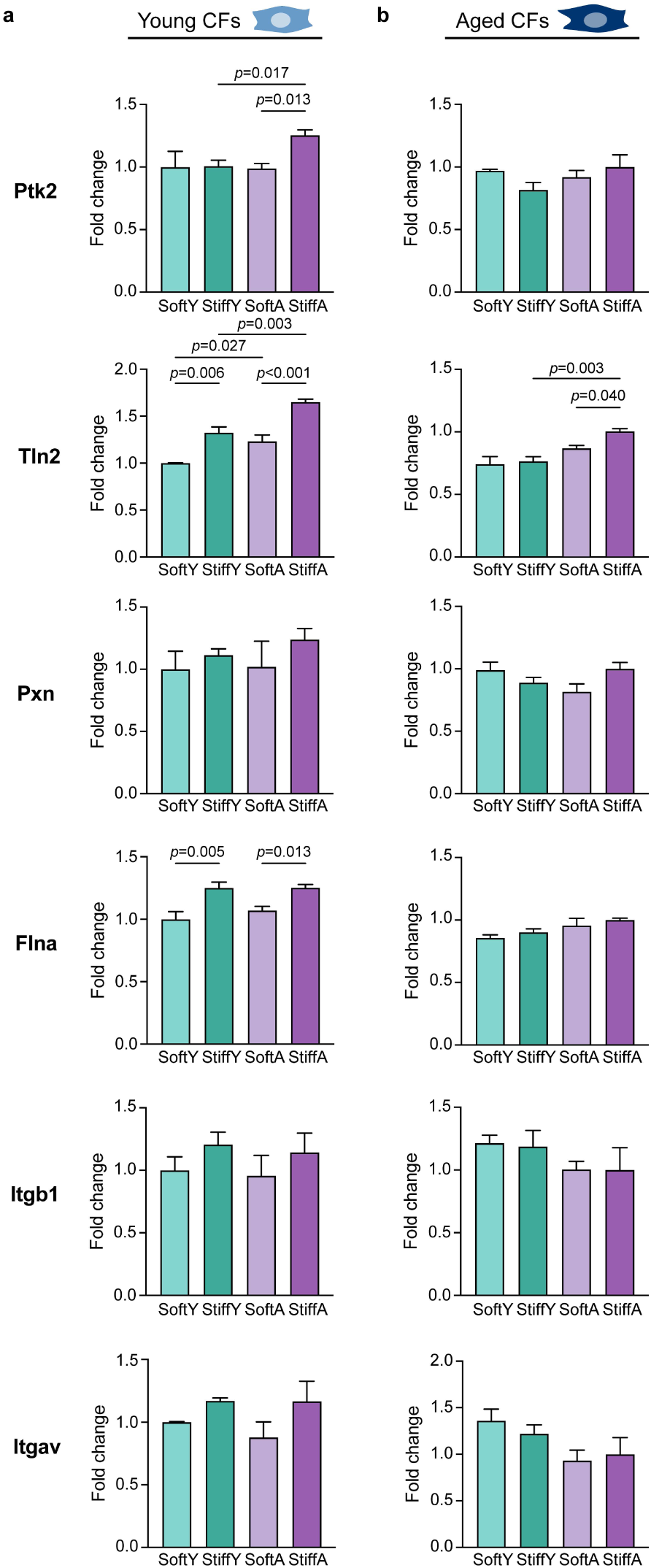
Expression of integrin and focal adhesion genes on DECIPHER scaffolds. Quantified RNA-seq data for protein tyrosine kinase 2 (Ptk2), talin 2 (Tln2), paxillin (Pxn), filamin A (Flna), integrin beta 1 (Itgb1), and integrin alpha v (Itgav) for YCFs (**a**) and ACFs (**b**) on DECIPHER samples. Normalized gene expression on different DECIPHER samples are colored as follows: SoftY - light green, StiffY - dark green, SoftA - light purple, StiffA - dark purple.

**Extended Data Fig. 10.**
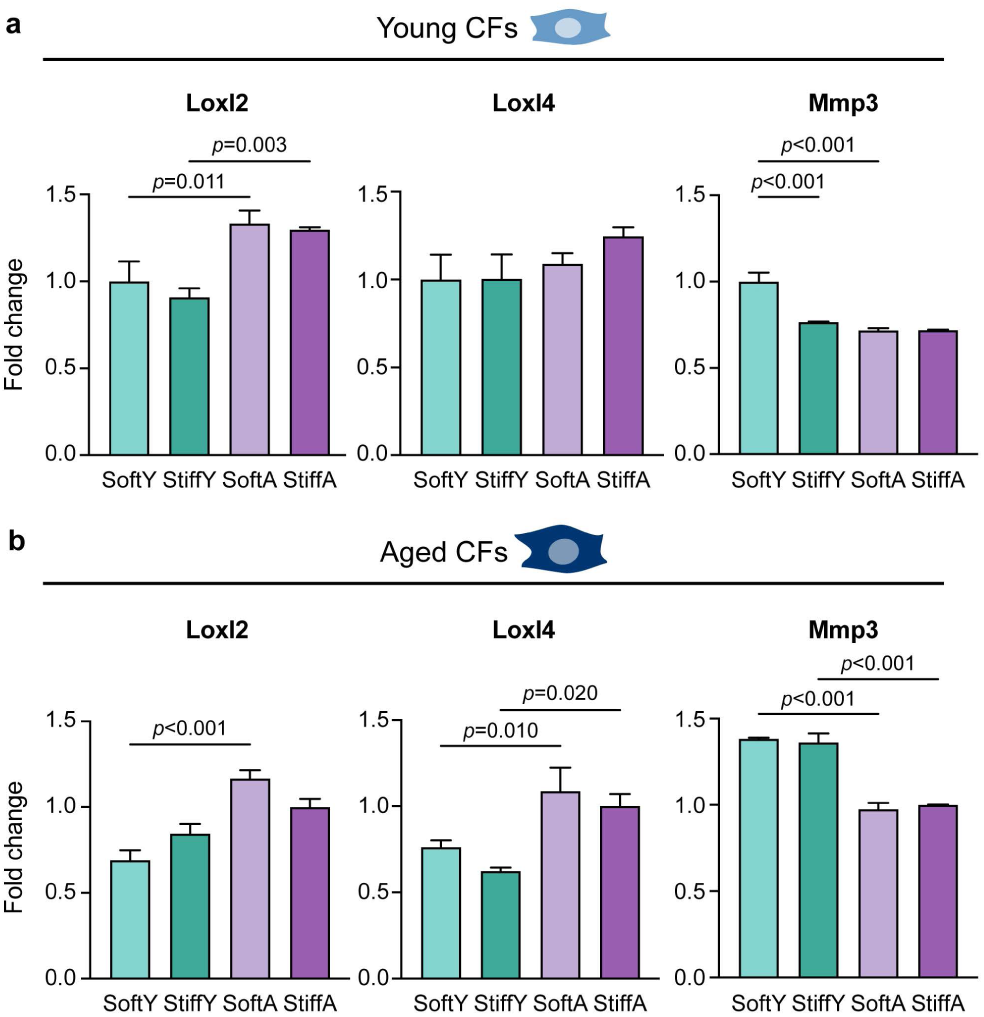
Matrix remodeling ability of CFs on DECIPHER scaffolds. Quantified RNA-seq data for genes encoding ECM crosslinking enzymes (LOXL2, LOXL4) and degrading enzyme (MMP3) expressed in YCFs (**a**) and ACFs (**b**). Normalized gene expression on different DECIPHER samples are colored as follows: SoftY - light green, StiffY - dark green, SoftA - light purple, StiffA - dark purple.

**Extended Data Fig. 11.**
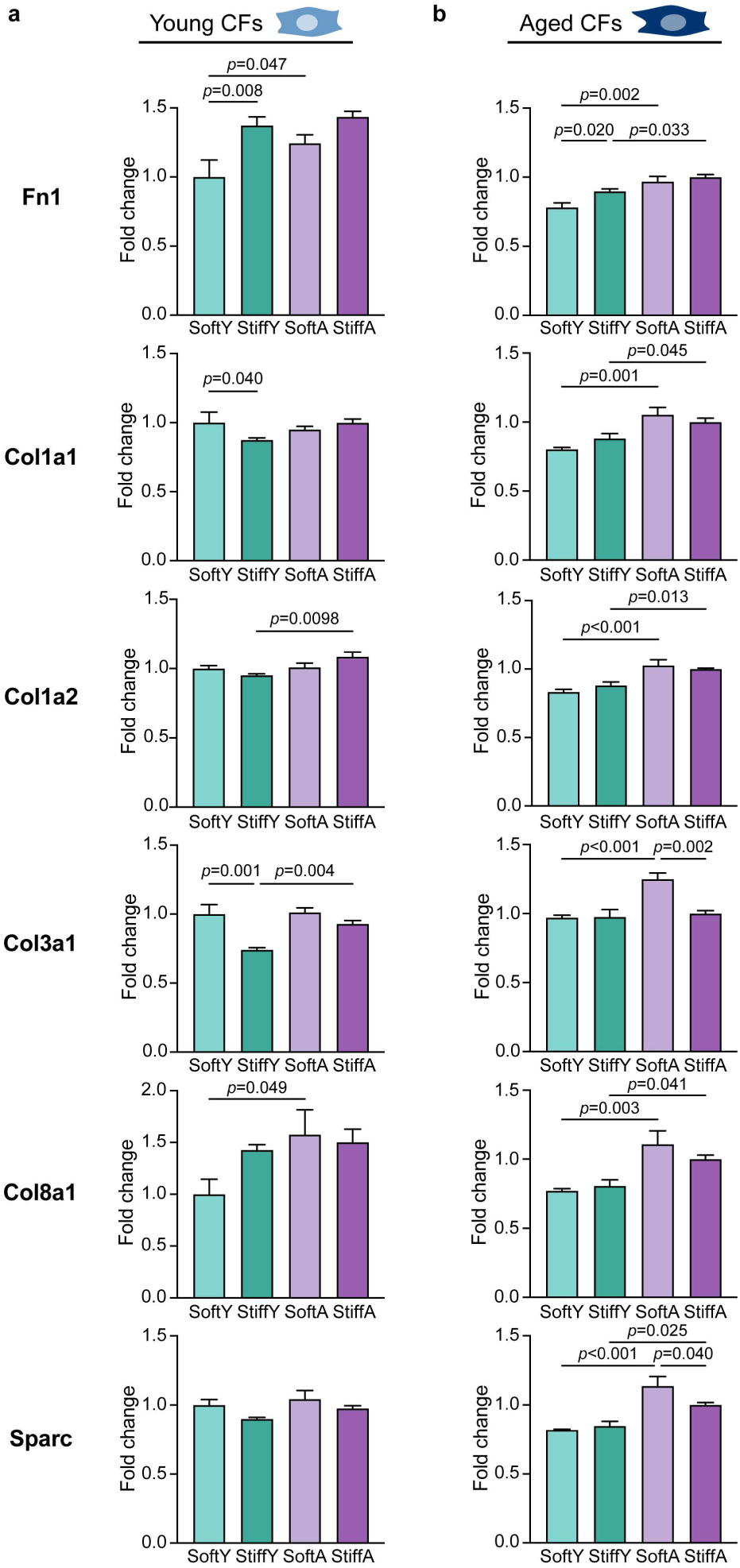
Synthesis of ECM components on DECIPHER scaffolds. Quantified RNA-seq data for fibronectin (Fn1); collagen I (Col1a1, Col1a2), III (Col3a1), VIII (Col8a1); and SPARC (Sparc) expressed in YCFs (**a**) and ACFs (**b**). Normalized gene expression on different DECIPHER samples are colored as follows: SoftY - light green, StiffY - dark green, SoftA - light purple, StiffA - dark purple.

**Extended Data Fig. 12.**
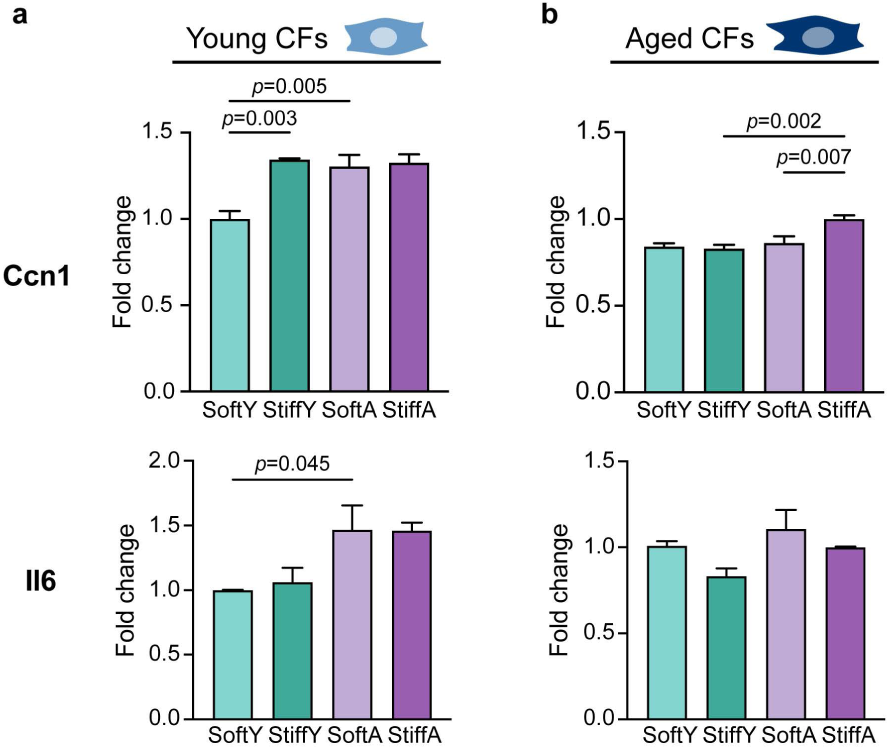
SASP expression on DECIPHER scaffolds. Quantified RNA-seq data for interleukin 6 (Il-6) and cellular communication network factor 1 (ccn1) YCFs (**a**) and ACFs (**b**) on DECIPHER samples. Normalized gene expression on different DECIPHER samples are colored as follows: SoftY - light green, StiffY - dark green, SoftA - light purple, StiffA - dark purple.

## Supplemental tables

**Table 1.** PA hydrogel stabilization solutions to mimic young/aged cardiac stiffness.

|  | **Acrylamide**<br>**%** | **Bis-**<br>**acrylamide**<br>**%** | **Irgacure**<br>**%** | **Formaldehyde**<br>**%** |
| --- | --- | --- | --- | --- |
| **Soft**<br>(Young heart mimic) | 10% | 0.15% | 0.5% | 0.4% |
| **Stiff**<br>(Aged heart mimic) | 12% | 0.25% | 0.5% | 0.4% |

**Table 2.**
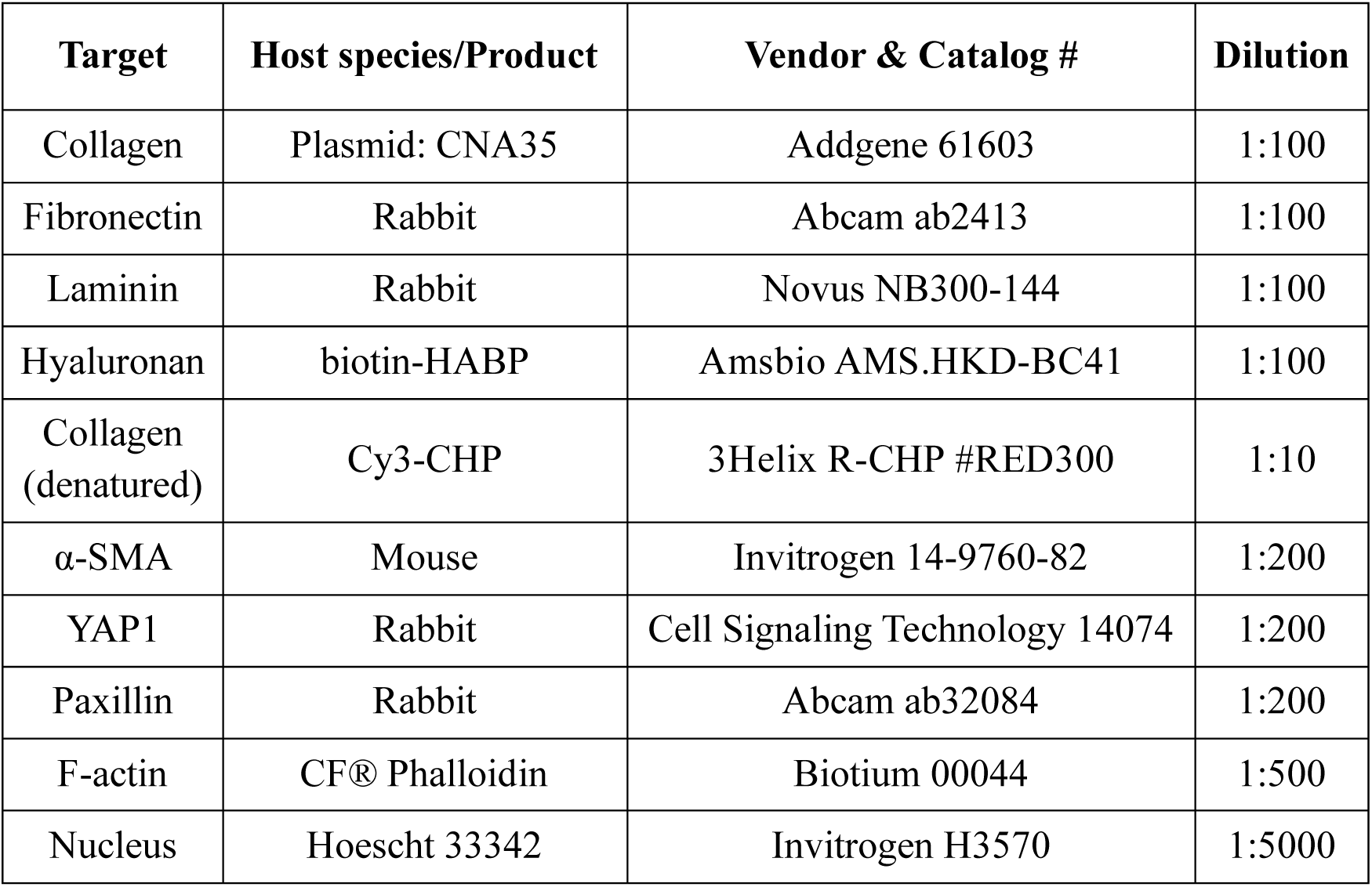
Antibodies and dyes used for IHC and ICC.

## References

1. Nance, M. E. et al. Attenuated sarcomere lengthening of the aged murine left ventricle observed using two-photon fluorescence microscopy. Am. J. Physiol. Circ. Physiol. 309, H918–H925 (2015).

2. Ozcebe, S. G., Bahcecioglu, G., Yue, X. S. & Zorlutuna, P. Effect of cellular and ECM aging on human iPSC-derived cardiomyocyte performance, maturity and senescence. Biomaterials 268, 120554 (2021).

3. Meschiari, C. A., Ero, O. K., Pan, H., Finkel, T. & Lindsey, M. L. The impact of aging on cardiac extracellular matrix. Geroscience 39, 7–18 (2017).

4. Antia, M., Baneyx, G., Kubow, K. E. & Vogel, V. Fibronectin in aging extracellular matrix fibrils is progressively unfolded by cells and elicits an enhanced rigidity response. Faraday Discuss 139, 229–249 (2008).

5. Yokota, T. et al. Type V Collagen in Scar Tissue Regulates the Size of Scar after Heart Injury. Cell 182, 545–562 (2020).

6. Herum, K. M., Choppe, J., Kumar, A., Engler, A. J. & McCulloch, A. D. Mechanical regulation of cardiac fibroblast profibrotic phenotypes. Mol Biol Cell 28, 1871–1882 (2017).

7. Wang, Z. et al. Snake venom-defined fibrin architecture dictates fibroblast survival and differentiation. Nat Commun 14, 1029 (2023).

8. Tomasek, J. J., Gabbiani, G., Hinz, B., Chaponnier, C. & Brown, R. A. Myofibroblasts and mechano-regulation of connective tissue remodelling. Nat Rev Mol Cell Bio 3, 349–363 (2002).

9. Talman, V. & Ruskoaho, H. Cardiac fibrosis in myocardial infarction—from repair and remodeling to regeneration. Cell Tissue Res 365, 563–581 (2016).

10. Angelini, A., Trial, J., Ortiz-Urbina, J. & Cieslik, K. A. Mechanosensing dysregulation in the fibroblast: a hallmark of the aging heart. Ageing Res Rev 63, 101150 (2020).

11. Weber, K. T., Sun, Y., Bhattacharya, S. K., Ahokas, R. A. & Gerling, I. C. Myofibroblast-mediated mechanisms of pathological remodelling of the heart. Nat Rev Cardiol 10, 15–26 (2013).

12. Cieslik, K. A., Trial, J., Carlson, S., Taffet, G. E. & Entman, M. L. Aberrant differentiation of fibroblast progenitors contributes to fibrosis in the aged murine heart: role of elevated circulating insulin levels. Faseb J 27, 1761–1771 (2013).

13. Chen, Z. et al. Intrafibrillar Crosslinking Enables Decoupling of Mechanical Properties and Structure of a Composite Fibrous Hydrogel. Adv. Mater. e2305964 (2023) doi:10.1002/adma.202305964.

14. Cunha, C. B. da et al. Influence of the stiffness of three-dimensional alginate/collagen-I interpenetrating networks on fibroblast biology. Biomaterials 35, 8927–8936 (2014).

15. Berger, A. J., Linsmeier, K. M., Kreeger, P. K. & Masters, K. S. Decoupling the effects of stiffness and fiber density on cellular behaviors via an interpenetrating network of gelatin-methacrylate and collagen. Biomaterials 141, 125–135 (2017).

16. Chung, K. et al. Structural and molecular interrogation of intact biological systems. Nature 497, 332–337 (2013).

17. Hwang, J. et al. Molecular assessment of collagen denaturation in decellularized tissues using a collagen hybridizing peptide. Acta Biomater 53, 268–278 (2017).

18. Aper, S. J. A. et al. Colorful Protein-Based Fluorescent Probes for Collagen Imaging. Plos One 9, e114983 (2014).

19. Ackers-Johnson, M. & Foo, R. S. Langendorff-Free Isolation and Propagation of Adult Mouse Cardiomyocytes. Methods Mol. Biol. 1940, 193–204 (2019).

20. Wershof, E. et al. A FIJI macro for quantifying pattern in extracellular matrix. Life Sci Alliance 4, e202000880 (2021).

21. Love, M. I., Huber, W. & Anders, S. Moderated estimation of fold change and dispersion for RNA-seq data with DESeq2. Genome Biol. 15, 550 (2014).

22. Wu, T. et al. clusterProfiler 4.0: A universal enrichment tool for interpreting omics data. Innov. 2, 100141 (2021).

23. Yu, G., Wang, L.-G., Han, Y. & He, Q.-Y. clusterProfiler: an R Package for Comparing Biological Themes Among Gene Clusters. *OMICS: A J*. Integr. Biol. 16, 284–287 (2012).

24. Chen, C. et al. TBtools: An Integrative Toolkit Developed for Interactive Analyses of Big Biological Data. Mol Plant 13, 1194–1202 (2020).

25. Tse, J. R. & Engler, A. J. Preparation of Hydrogel Substrates with Tunable Mechanical Properties. Curr Protoc Cell Biology 47, 10.16.1–10.16.16 (2010).

26. Crapo, P. M., Gilbert, T. W. & Badylak, S. F. An overview of tissue and whole organ decellularization processes. Biomaterials 32, 3233–3243 (2011).

27. Nakayama, K. H., Batchelder, C. A., Lee, C. I. & Tarantal, A. F. Decellularized Rhesus Monkey Kidney as a Three-Dimensional Scaffold for Renal Tissue Engineering. Tissue Eng. Part A 16, 2207–2216 (2010).

28. Fathi, I. et al. Decellularized Whole-Organ Pre-vascularization: A Novel Approach for Organogenesis. Front. Bioeng. Biotechnol. 9, 756755 (2021).

29. Jeong, D. et al. Matricellular Protein CCN5 Reverses Established Cardiac Fibrosis. J. Am. Coll. Cardiol. 67, 1556–1568 (2016).

30. Chen, M. S., Lee, R. T. & Garbern, J. C. Senescence mechanisms and targets in the heart. Cardiovasc Res 118, 1173–1187 (2021).

31. Pesce, M. et al. Cardiac fibroblasts and mechanosensation in heart development, health and disease. Nat Rev Cardiol 1–16 (2022) doi:10.1038/s41569-022-00799-2.

32. Ma, Y., Iyer, R. P., Jung, M., Czubryt, M. P. & Lindsey, M. L. Cardiac Fibroblast Activation Post-Myocardial Infarction: Current Knowledge Gaps. Trends Pharmacol Sci 38, 448–458 (2017).

33. Wang, J., Liu, S., Heallen, T. & Martin, J. F. The Hippo pathway in the heart: pivotal roles in development, disease, and regeneration. Nat. Rev. Cardiol. 15, 672– 684 (2018).

34. Kanchanawong, P. & Calderwood, D. A. Organization, dynamics and mechanoregulation of integrin-mediated cell–ECM adhesions. Nat Rev Mol Cell Bio 1–20 (2022) doi:10.1038/s41580-022-00531-5.

35. Devarasou, S., Kang, M., Kwon, T. Y., Cho, Y. & Shin, J. H. Fibrous Matrix Architecture-Dependent Activation of Fibroblasts with a Cancer-Associated Fibroblast-like Phenotype. ACS Biomater. Sci. Eng. 9, 280–291 (2023).

36. DeLeon-Pennell, K. Y., Meschiari, C. A., Jung, M. & Lindsey, M. L. Chapter Two Matrix Metalloproteinases in Myocardial Infarction and Heart Failure. Prog. Mol. Biol. Transl. Sci. 147, 75–100 (2017).

37. Mizuno, T., Yau, T. M., Weisel, R. D., Kiani, C. G. & Li, R.-K. Elastin Stabilizes an Infarct and Preserves Ventricular Function. Circulation 112, I81–8 (2005).

38. Meyer, K., Hodwin, B., Ramanujam, D., Engelhardt, S. & Sarikas, A. Essential Role for Premature Senescence of Myofibroblasts in Myocardial Fibrosis. J. Am. Coll. Cardiol. 67, 2018–2028 (2016).

39. Zhu, F. et al. Senescent Cardiac Fibroblast Is Critical for Cardiac Fibrosis after Myocardial Infarction. Plos One 8, e74535 (2013).

40. Coppé, J.-P., Desprez, P.-Y., Krtolica, A. & Campisi, J. The Senescence-Associated Secretory Phenotype: The Dark Side of Tumor Suppression. Pathol.: Mech. Dis. 5, 99–118 (2010).

41. Basara, G., Ozcebe, S. G., Ellis, B. W. & Zorlutuna, P. Tunable Human Myocardium Derived Decellularized Extracellular Matrix for 3D Bioprinting and Cardiac Tissue Engineering. Prog Coll Pol Sci S 7, 70 (2021).

42. Kim, H. et al. Light-Activated Decellularized Extracellular Matrix-Based Bioinks for Volumetric Tissue Analogs at the Centimeter Scale. Adv Funct Mater 31, 2011252 (2021).

43. Spang, M. T. et al. Intravascularly infused extracellular matrix as a biomaterial for targeting and treating inflamed tissues. *Nat*. Biomed. Eng. 7, 94–109 (2023).

44. Umehara, T. et al. Female reproductive life span is extended by targeted removal of fibrotic collagen from the mouse ovary. Sci Adv 8, eabn4564 (2022).

45. Cho, S., Discher, D. E., Leong, K. W., Vunjak-Novakovic, G. & Wu, J. C. Challenges and opportunities for the next generation of cardiovascular tissue engineering. Nat Methods 1–8 (2022) doi:10.1038/s41592-022-01591-3.

46. Engler, A. J., Sen, S., Sweeney, H. L. & Discher, D. E. Matrix Elasticity Directs Stem Cell Lineage Specification. Cell 126, 677–689 (2006).

47. Hadden, W. J. et al. Stem cell migration and mechanotransduction on linear stiffness gradient hydrogels. Proceedings of the National Academy of Sciences 114, 5647–5652 (2017).

48. Iskratsch, T., Wolfenson, H. & Sheetz, M. P. Appreciating force and shape — the rise of mechanotransduction in cell biology. Nat. Rev. Mol. Cell Biol. 15, 825–833 (2014).

49. Russo, J. D. et al. Integrin α5β1 nano-presentation regulates collective keratinocyte migration independent of substrate rigidity. eLife 10, e69861 (2021).

50. Trappmann, B. et al. Extracellular-matrix tethering regulates stem-cell fate. Nat. Mater. 11, 642–649 (2012).

51. Young, J. L. & Engler, A. J. Hydrogels with time-dependent material properties enhance cardiomyocyte differentiation in vitro. Biomaterials 32, 1002–1009 (2011).

52. Walker, C. J. et al. Nuclear mechanosensing drives chromatin remodelling in persistently activated fibroblasts. *Nat*. Biomed. Eng. 5, 1485–1499 (2021).

